# Alzheimer’s disease-associated Presenilin 2 N141I mutation impairs neuronal lipid homeostasis and mitochondrial dynamics through selective downregulation of the Golgi exchange factor *Gbf1*

**DOI:** 10.64898/2026.05.20.726466

**Authors:** Shirine Saleki, Claire Wabant, Axelle Loriot, Serena Stanga, Julien Masquelier, Giulio G Muccioli, Nuria Suelves, Pascal Kienlen-Campard

**Affiliations:** Aging and Dementia lab, Institute of Neuroscience, UCLouvain, Brussels, Belgium; Group of Computational Biology and Bioinformatics, de Duve Institute, UCLouvain, Brussels, Belgium; Neuroscience Institute Cavalieri Ottolenghi, Department of Neuroscience Rita Levi Montalcini, University of Turin, Turin, Italy; Bioanalysis and Pharmacology of Bioactive Lipids, Louvain Drug Research Institute, UCLouvain, Brussels, Belgium; Avenue Mounier 53 bte B1.53.02, 1200 Brussels, Belgium

**Keywords:** Alzheimer’s disease, presenilins, lipid homeostasis, mitochondria, neuron

## Abstract

Presenilin 2 (PS2) mutations cause familial Alzheimer’s disease, yet their effects beyond amyloid processing remain poorly understood. Here, we investigated how PS2 deletion and the N141I mutation affect neuronal lipid homeostasis and mitochondrial dynamics in mouse primary neurons. Both PS2 deletion and N141I mutation reduced neuronal lipid content. However, exogenous lipid supplementation rescued this deficit only in N141I-expressing neurons, indicating a partial loss-of-function effect. N141I neurons also displayed reduced OPA1, a mitochondrial fusion regulator, restored by lipid supplementation. RNA-sequencing identified *Gbf1*, a Golgi-specific guanine nucleotide exchange factor, as selectively downregulated in N141I but not knockout tissue, which was confirmed at the protein level in mouse brain and primary neurons. *Gbf1* knockdown in mouse embryonic fibroblasts (MEFs) recapitulated the N141I lipid profile. Together, these findings reveal a PS2-GBF1-lipid-mitochondria axis disrupted specifically by the N141I mutation, suggesting an amyloid-independent pathway contributing to neurodegeneration and identifying potential therapeutic targets for familial Alzheimer’s disease.

## INTRODUCTION

Alzheimer’s disease (AD) is the leading cause of dementia worldwide, clinically characterized by progressive cognitive decline and neuropathologically defined by the accumulation of extracellular amyloid-β (Aβ) plaques and intracellular neurofibrillary tau tangles. Aβ is generated by the sequential proteolytic processing of the amyloid precursor protein (APP) by the β- and γ-secretase complexes, the latter requiring either presenilin 1 (PS1) or presenilin 2 (PS2) as its catalytic subunit^1,2^. Mutations in the genes encoding PS1 (*PSEN1*) and PS2 (*PSEN2*), together with *APP* mutations, account for the majority of familial AD (FAD) cases^3–5^. While the amyloid cascade hypothesis has dominated AD research for decades^1,6^, an increasing body of evidence points to cellular processes including autophagy defects^7^, mitochondrial dysfunction, and possibly associated lipid dyshomeostasis, as early, mechanistically central features of the disease^8,9^. Whether these features precede and drive amyloid pathology, occur independently, or arise as a simple consequence of it, remains to be fully elucidated. In particular, lipid dyshomeostasis in an AD context needs to be further characterized in order to develop lipidomics-driven strategies and evaluate the potential of lipid-based therapeutic approaches^10^.

The brain is an exceptionally lipid-rich organ. Lipids constitute more than 50% of its dry weight, with phospholipids, sphingolipids, cholesterol, and glycolipids collectively governing membrane structure and fluidity, and the organization of specialized microdomains such as lipid rafts^11^. Lipidomic and metabolomic studies have consistently shown early alterations in the levels of various lipid classes in AD brains, revealing multifaceted interactions between lipid metabolism and key AD pathogenic mechanisms, including amyloidogenesis, bioenergetic deficit, oxidative stress, neuroinflammation, and myelin degeneration^12^. Alterations in lipid composition are well documented in AD, characterized by reductions in phospholipids and sulfatides, along with increases in cholesterol, cholesteryl esters, and triglycerides^13^. Phospholipids play a pivotal role in the activity of the γ-secretase and APP processing, and investigations in AD patient-derived cell cultures have uncovered shifts in various phospholipid types, including phosphatidylcholine (PC), phosphatidylethanolamine (PE), and phosphatidylinositol (PI), along with dysregulation of the enzymes responsible for their metabolism^14^. Critically, pathological presenilin mutations (i.e. *PSEN1*) themselves result in increased cholesterol and decreased sphingomyelin levels^15^, placing presenilins at the interface of both amyloid processing and lipid homeostasis. Consistent with this, membrane fluidity and cholesterol distribution in detergent-resistant membrane fractions are altered in presenilin-deficient mouse embryonic fibroblasts^16,17^ and FAD-linked mutations in both *PSEN1* and *PSEN2* cause an imbalance in phosphatidylinositol 4,5-bisphosphate (PIP_2_) metabolism^18^.

Mitochondrial dysfunction constitutes a parallel hallmark of AD, emerging early and correlating with cognitive decline^19^. Mitochondrial dysfunction and oxidative stress are frequently observed in early stages of AD, and a mitochondrial cascade hypothesis has been advanced in which mitochondrial impairment, including abnormal morphology, disrupted oxidative phosphorylation (OXPHOS), increased reactive oxygen species (ROS) production, and impaired biogenesis, may represent early pathological events driving downstream amyloid and tau accumulation^20^. The endoplasmic reticulum (ER)-mitochondria interface, known as mitochondria-associated membranes (MAMs), is a key hub linking lipid metabolism, calcium signaling, and mitochondrial function. While both PS1 and PS2 are enriched at MAMs^21^, PS2 specifically regulates ER-mitochondria coupling and calcium crosstalk^22,23^. MAMs also act as lipid raft-like domains that coordinate the synthesis and transfer of phospholipids and cholesterol to mitochondria, thereby shaping membrane composition, cristae architecture, and metabolic enzyme activity^24,25^. In AD, APP and presenilin mutants accumulate aberrantly in MAMs and disrupt both calcium homeostasis and lipid transfer, promoting neurotoxic Aβ generation. The convergence of lipid dysregulation and mitochondrial impairment at the MAMs thus suggests these two pathological processes are not independent but are mechanistically coupled, and that disruption of this coupling could be a key event in AD neurodegeneration.

Among the presenilins, PS2 occupies a particularly compelling position. While PS1 predominantly localizes at the plasma membrane and carries the majority of known FAD mutations^26^, PS2-dependent γ-secretase complexes are predominantly targeted to late endosomes and lysosomes^27^ – compartments associated with the production of a more pathogenic intracellular Aβ pool^28^. PS2 FAD mutations, including the N141I substitution (the first pathogenic mutation described on *PSEN2*^29,30^), strongly increase ER-mitochondria interaction and mitochondrial calcium uptake^24,31,32^, and have been associated with altered γ-secretase activity^4,33^, disrupted calcium signaling^34^, and neuronal death^27,35^. The broader mechanistic significance of PS2’s γ-secretase-independent functions, including calcium homeostasis, mitochondrial function, autophagy, and lipid metabolism, remains incompletely understood. It is, however, increasingly recognized that these non-catalytic roles may be as critical to AD pathogenesis as Aβ production itself^20^.

We have previously demonstrated, in mouse embryonic fibroblasts (MEFs) derived from PS2 knockout (PS2KO) and PS1/PS2 double knockout (PSdKO) mice that the absence of PS2 – but not PS1 – leads to significant impairment in mitochondrial OXPHOS capacity. This results in decreased basal respiration, coupling efficiency, and spare respiratory capacity, alongside structural defects in mitochondrial cristae and a decreased NAD^+^/NADH ratio. These functional deficits were compensated by an upregulation of glycolytic flux and were fully rescued by stable re-expression of PS2, demonstrating their specific PS2 dependence^36^.

Building on this basis, we investigated PS2-dependent molecular mechanisms in neurons, a cell type relevant to AD pathogenesis where the consequences of PS2 dysfunction may be particularly pronounced, given their high oxidative demands and marked reliance on lipid-rich membranes for synaptic function. We employed a series of complementary genetic models, including PS2 homozygous knockout (PS2-/-) and heterozygous knockout (PS2+/-) mice, a transgenic model expressing the FAD-linked N141I mutation on a PS2+/-background (PS2N141I mice)^33^, and corresponding primary neuronal cultures, to dissect whether the N141I mutation mimics or diverges from the loss of PS2 function, with a specific focus on neuronal lipid content. We next aimed at identifying PS2-dependent downstream targets that could relate the PS2-dependent phenotype to lipid homeostasis and mitochondrial function. Our findings reveal a partial loss-of-function effect of the N141I mutation on PS2’s role in lipid metabolism, demonstrate a lipid-dependent mitochondrial phenotype in PS2N141I neurons, and identify *Gbf1* (Golgi-specific brefeldin A-resistance guanine nucleotide exchange factor 1) as a novel, mutation-specific downstream target of PS2, collectively pointing to a PS2-GBF1-lipid-mitochondria axis as a potentially important, Aβ-independent pathway in AD pathogenesis.

## RESULTS

### Presenilin 2 deletion and the N141I mutation each reduce lipid content in primary neurons

To obtain an unbiased assessment of PS2-dependent lipid alterations, we first performed lipidomic analyses in mouse embryonic fibroblasts (MEFs) lacking PS1 (PS1-/-) or PS2 (PS2-/-). These analyses revealed significantly decreased levels of sphingomyelin, ceramide, and phosphatidylinositol (PI) in PS2-/- but not PS1-/- MEFs (Figure S1), highlighting a PS2-dependent lipid profile alteration. Based on this observation, we next investigated whether similar lipid changes occur in a neuronal context.

PS1 and PS2 protein levels were first assessed by Western blotting in primary neurons derived from WT, PS2-/-, PS2+/-, and PS2N141I embryos, to verify the expression status of PS2 across all genetic models and confirm the absence of compensatory changes in PS1 levels. As expected, PS2 was undetectable in PS2-/- neurons, confirming complete loss of the endogenous protein, and no significant compensatory changes were observed on PS1 levels (Figure S2A). As reported before, the effect of the PS2 N141I mutation was studied in a PS2+/- background to limit the global overexpression of PS2^33^. In PS2N141I neurons, PS2 levels were logically elevated relative to PS2+/- controls, consistent with expression of the human *PSEN2* N141I transgene in addition to the remaining endogenous allele, and PS1 levels were unaltered (Figure S2B).

To complement the transgenic approach and control for potential confounds related to the PS2N141I transgenic background, WT primary neurons were infected with lentiviral vectors expressing either wild-type human PS2 (LV-PSEN2WT) or PS2 carrying the N141I mutation (LV-PSEN2N141I). Western blotting confirmed comparable PS2 protein levels between the two infected conditions (Figure S2C), ensuring that subsequent comparisons using the lentiviral system reflect the effect of the N141I mutation rather than differences in PS2 expression levels.

Lipid content was assessed by PhenoVue™ Nile Red fluorescent staining in combination with MAP2 (Microtubule-Associated Protein 2) immunostaining to identify neurons. Nile Red is a solvatochromic dye that partitions into lipid-rich environments and emits at distinct wavelengths depending on lipid polarity, allowing separate quantification of neutral lipids (principally triglycerides and cholesterol esters) and phospholipids^37^. Relative fluorescence intensity (RFU) was normalized to Hoechst signal to correct for cell density.

In PS2-/- neurons, neutral lipid content was unchanged relative to WT neurons, whereas phospholipid fluorescence was significantly reduced (Figure 1A), suggesting a selective effect in phospholipid homeostasis.

**Figure 1.**
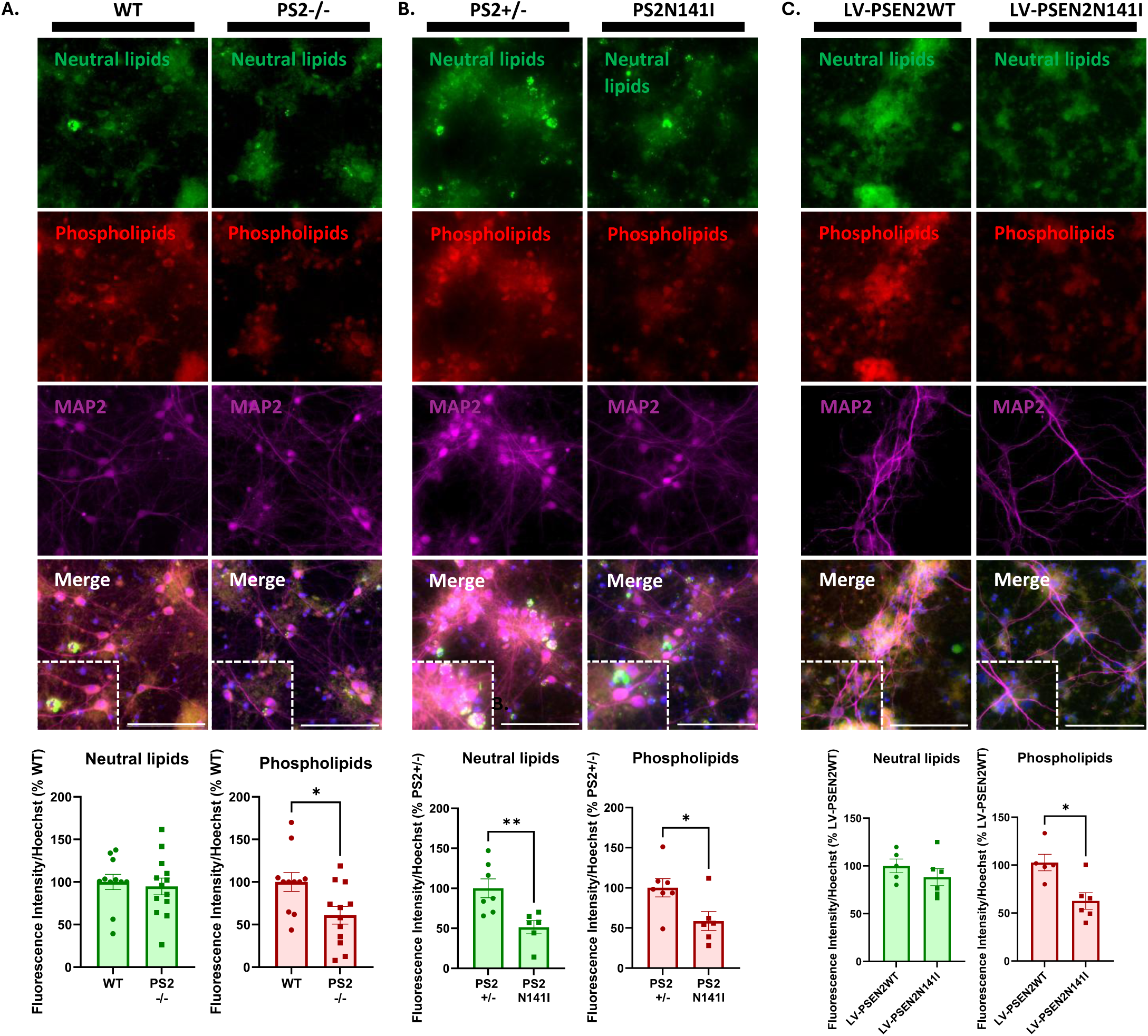
Presenilin 2 deletion and the N141I mutation reduce neuronal lipid content. (A) PhenoVue Nile Red staining combined with MAP2 immunostaining of WT and PS2-/- primary neurons. Inlets show zoomed-in images. Relative fluorescence intensity (RFU) for Neutral lipids (in green) or Phospholipids (in red) was calculated and normalized to Hoechst RFU. The levels in the control group (WT) were set as 100%. \**p* < 0.05 (Unpaired t-test, N = 5 cultures, n = 11-13 embryos/group). (B) PhenoVue Nile Red staining combined with MAP2 immunostaining of PS2+/- and PS2N141I primary neurons. Inlets show zoomed-in images. Relative fluorescence intensity (RFU) for Neutral lipids (in green) or Phospholipids (in red) was calculated and normalized to Hoechst RFU. The levels in the control group (PS2+/-) were set as 100%. \**p* < 0.05, \*\**p* < 0.01 (Unpaired t-test, N = 3 cultures, n = 6-7 embryos/group). (C) PhenoVue Nile Red staining combined with MAP2 immunostaining of PS2+/- primary neurons infected with a lentivirus (LV) expressing PS2 WT (LV-PSEN2WT) or PS2 N141I (LV-PSEN2N141I). Inlets show zoomed-in images. Relative fluorescence intensity (RFU) for Neutral lipids (in green) or Phospholipids (in red) was calculated and normalized to Hoechst RFU. The levels in the control group (LV-PSEN2WT) were set as 100%. \**p* < 0.05 (Unpaired t-test, N = 3 cultures, n = 5-6 embryos/group). Scale bar: 100 µm. Data are presented as mean ± SEM.

In transgenic PS2N141I neurons, both neutral lipid and phospholipid content were significantly decreased compared to PS2+/- controls (Figure 1B). The broader lipid deficit observed with the N141I mutation – affecting both lipid classes compared to only phospholipids in the knockout – suggests that the mutation does not simply recapitulate the loss-of-function phenotype but rather exerts additional effects on lipid metabolism.

In the lentiviral model, neutral lipid content was unchanged between LV-PSEN2N141I and LV-PSEN2WT neurons, whereas a significantly decreased phospholipid content was observed in the LV-PSEN2N141I condition (Figure 1C). This more selective phospholipid deficit, in contrast to the broader phenotype seen in the transgenic model, may reflect quantitative differences in PS2 expression levels between the two systems. Although both models express N141I on a PS2+/- background, the overall PS2 expression level (endogenous allele and transgene or lentiviral vector) may differ, and the transgenic model may achieve higher total PS2 N141I levels that are sufficient to additionally perturb neutral lipid metabolism. Importantly, the phospholipid deficit is replicated across all three models and represents the most consistent lipid phenotype associated with the N141I mutation.

Collectively, these data establish that both complete PS2 loss and the N141I FAD mutation impair neuronal lipid homeostasis. However, while the transgenic model shows deficits across neutral lipids and phospholipids, the lentiviral PSEN2N141I overexpression model and the PS2 knockout model reveal a more restricted effect limited to phospholipids, highlighting the importance of the expression context in shaping the lipid phenotype.

### Exogenous lipid supplementation rescues the lipid deficit in Presenilin 2-mutant but not in Presenilin 2-deficient neurons

Standard neuronal culture medium (Neurobasal®) is serum-free and lipid-deprived by design. To test whether the lipid deficits observed in each model could be restored by providing an exogenous lipid source, neurons were supplemented with a chemically defined Gibco™ Lipid Concentrate (LC; 1:100 dilution) – described as a combination of unsaturated and saturated fatty acids – for 72 hours prior to lipid evaluation. Lipid content was again assessed by PhenoVue™ Nile Red staining.

In PS2-/- neurons supplemented with the lipid concentrate, phospholipid content remained significantly lower than in WT neurons (Figure 2A), mirroring the deficit observed in the absence of exogenous lipids. Neutral lipid levels were also unchanged between WT and PS2-/- neurons under supplemented conditions. These results indicate that complete PS2 deficiency renders neurons unable to restore their lipid content even when exogenous lipids are available.

**Figure 2.**
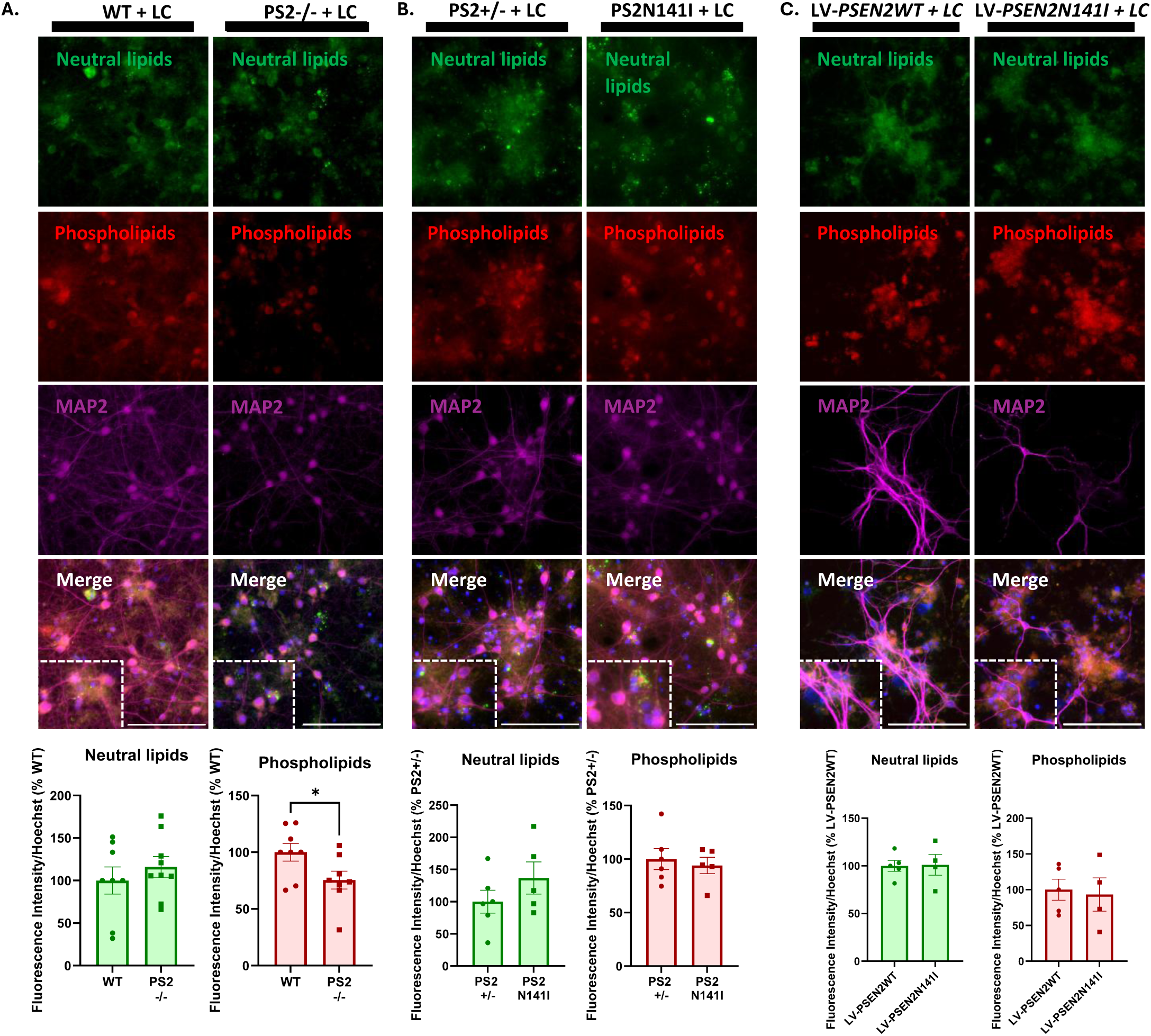
Exogenous lipid supplementation rescues the lipid deficit in PS2N141I but not PS2-/-primary neurons. (A) PhenoVue Nile Red staining combined with MAP2 immunostaining of WT and PS2-/- primary neurons cultivated in NB27 supplemented (+) with the Gibco™ lipid concentrate (LC). Inlets show zoomed-in images. Relative fluorescence intensity (RFU) for Neutral lipids (in green) or Phospholipids (in red) was calculated and normalized to Hoechst RFU. The levels in the control group (WT) were set as 100%. \**p* < 0.05 (Unpaired t-test, N = 4 cultures, n = 8-9 embryos/group) (B) PhenoVue Nile Red staining combined with MAP2 immunostaining of PS2+/- and PS2N141I primary neurons cultivated in NB27 supplemented (+) with the Gibco™ lipid concentrate. Inlets show zoomed-in images. Relative fluorescence intensity (RFU) for Neutral lipids (in green) or Phospholipids (in red) was calculated and normalized to Hoechst RFU. The levels in the control group (PS2+/-) were set as 100%. Non-significant (Unpaired t-test, N = 3 cultures, n = 5-6 embryos/group). (C) PhenoVue Nile Red staining combined with MAP2 immunostaining of PS2+/- primary neurons infected with a lentivirus (LV) expressing PS2 WT (LV-*PSEN2WT*) or PS2 N141I (LV-*PSEN2N141I*) cultivated in NB27 supplemented (+) with the Gibco™ lipid concentrate (LC). Inlets show zoomed-in images. Relative fluorescence intensity (RFU) for Neutral lipids (in green) or Phospholipids (in red) was calculated and normalized to Hoechst RFU. The levels in the control group (LV-*PSEN2WT*) were set as 100%. Non-significant (Unpaired t-test, N = 3 cultures, n = 4-5 embryos/group). Scale bar: 100 µm. Data are presented as mean ± SEM.

In contrast, in the PS2N141I transgenic model, supplementation with the lipid concentrate fully abolished the lipid deficit: no significant differences in either neutral lipid or phospholipid content were detected between PS2N141I and PS2+/- neurons after supplementation (Figure 2B). Consistent results were obtained in the lentiviral model: no significant differences in neutral lipid or phospholipid content were found between LV-PSEN2N141I and LV-PSEN2WT neurons following lipid supplementation (Figure 2C).

The divergent response to exogenous lipid supplementation between PS2-/- (non-rescued) and PS2N141I (rescued) neurons is a key finding. It demonstrates that the N141I mutation causes only a partial impairment of PS2’s lipid-related functions: while the mutation is sufficient to disrupt basal lipid homeostasis under lipid-deprived conditions, the reversal of the lipid deficit upon supplementation implies that the mutant protein, albeit dysfunctional, retains sufficient residual activity to permit exogenous lipid uptake or processing, though the specific step (uptake, intracellular trafficking, or esterification) has not been directly measured. This partial loss-of-function interpretation is consistent with previous evidence that PS2 N141I retains residual but altered γ-secretase activity^4,5^.

### The N141I mutation selectively impairs mitochondrial dynamics proteins in a lipid-dependent manner

Given that PS2 has been linked to mitochondrial homeostasis^22,31,36^ and that lipid homeostasis is tightly regulated by mitochondria, we next examined mitochondrial dynamics as a potential point of convergence. Mitochondrial dynamics, through the regulation of fusion and fission processes, plays an essential role in maintaining mitochondrial function, and was therefore assessed in primary neurons. We quantified the protein levels of three key regulators of mitochondrial morphology and dynamics by Western blotting: OPA1 (also called Dynamin-like 120 kDa protein), which mediates inner mitochondrial membrane (IMM) fusion and cristae organization; DRP1 (Dynamin-related Protein 1), which drives mitochondrial fission; and Mitofusin 2, which mediates outer mitochondrial membrane fusion^38^. These were assessed in WT vs. PS2-/- and PS2+/- vs. PS2N141I primary neurons, with or without lipid concentrate supplementation.

In PS2-/- neurons, no significant changes in OPA1, DRP1, or Mitofusin 2 protein levels were detected under either culture condition (Figure 3A). In PS2N141I neurons cultured without lipid supplementation, OPA1 levels were significantly reduced compared to PS2+/- neurons (Figure 3B). DRP1 and Mitofusin 2 levels also trended lower (DRP1: *p* = 0.1306; Mitofusin 2: *p* = 0.0813), but these differences did not reach statistical significance. When PS2N141I neurons were supplemented with the lipid concentrate, OPA1 levels were restored to PS2+/- levels, and the trends for DRP1 and Mitofusin 2 were no longer present, indicating that these mitochondrial phenotypes are fully reversed by lipid supplementation.

**Figure 3.**
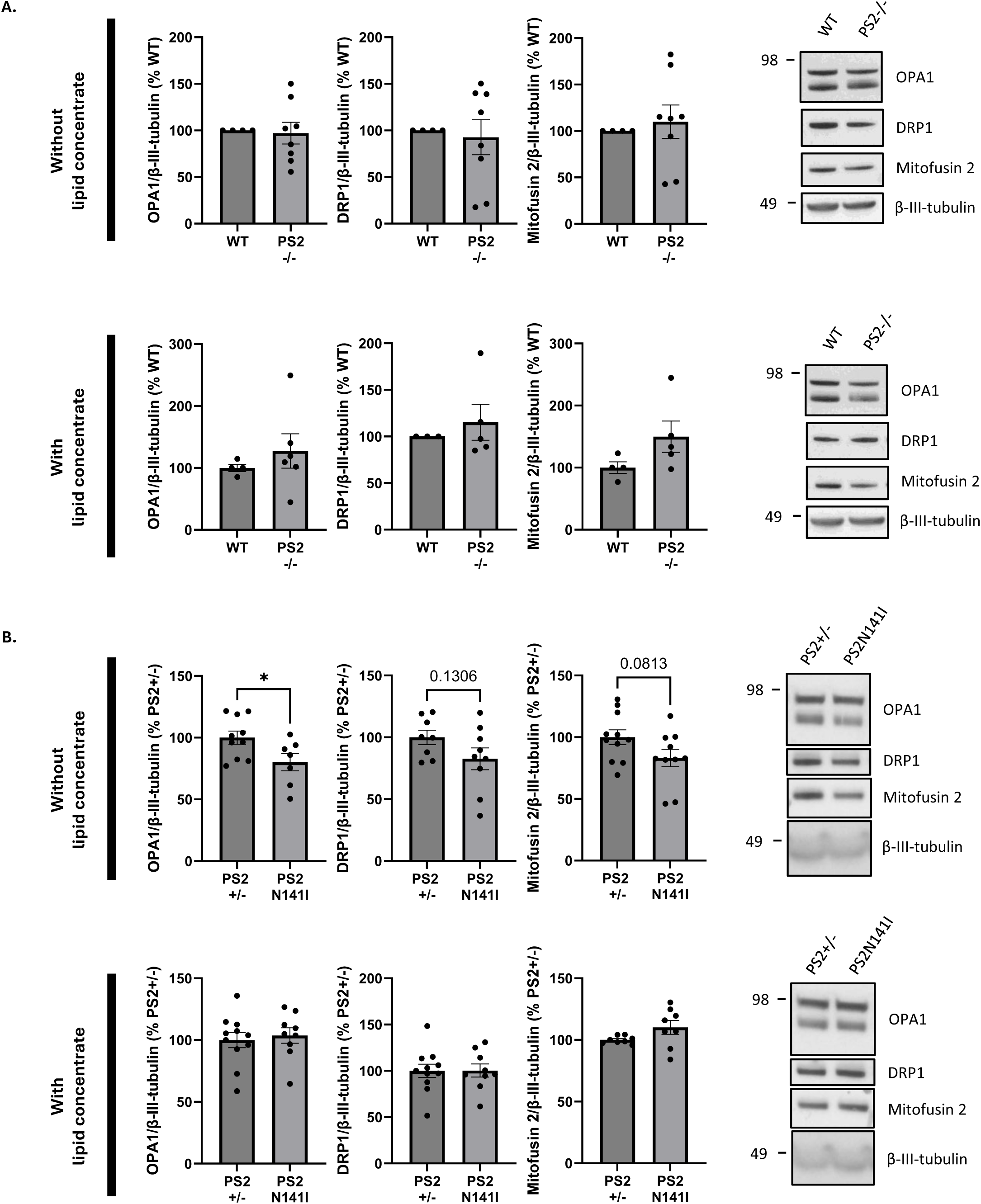
OPA1 levels are selectively reduced in PS2N141I neurons and rescued by exogenous lipid supplementation. (A) Measure of OPA1, DRP1 and Mitofusin 2 protein levels in WT and PS2-/- primary neurons supplemented or not with a lipid concentrate. β-III-tubulin was used as loading control, and the levels in the control group (WT) were set as 100%. Non-significant (Unpaired t-test, N = 4 experiments, n = 4-8 embryos/group). (B) Measure of OPA1, DRP1 and Mitofusin 2 protein levels in PS2+/- and PS2N141I primary neurons supplemented or not with a lipid concentrate. β-III-tubulin was used as loading control, and the levels in the control group (PS2+/-) were set as 100%. \**p* < 0.05 (Unpaired t-test, N = 4 experiments, n = 7-11 embryos/group). Data are presented as mean ± SEM.

The selective reduction of OPA1 in PS2N141I neurons and its rescue by exogenous lipids mirror the lipid disregulation pattern observed in the Nile Red lipid staining (Figure 1; Figure 2). However, the absence of mitochondrial protein dysregulation in PS2-/- neurons, despite the pronounced lipid deficit, suggests that a reduction in lipid content *per se* is not sufficient to drive mitochondrial dynamics alterations, and that the mitochondrial phenotype observed in PS2N141I neurons may instead reflect a qualitative change in lipid composition or an additional pathological gain of function conferred by the N141I mutation. The non-significant trends for DRP1 (*p* = 0.1306) and Mitofusin 2 (*p* = 0.0813) are noted but, given the underpowered sample size (N = 4 experiments), do not support further interpretation at this stage. Importantly, OPA1, DRP1, and Mitofusin 2 levels are used as protein-level proxies for mitochondrial dynamics pending mitochondrial morphology validation by direct imaging.

### Golgi exchange factor 1 Gbf1 is identified as a N141I mutation-specific downstream target responsible for Presenilin 2-dependent phospholipid deficit

To identify molecular mediators of the PS2-dependent lipid phenotype, RNA-sequencing was performed on hippocampal extracts from WT, PS2+/-, PS2-/-, and PS2N141I mice at 3 months of age (n = 4 per genotype). The top 10 differentially expressed genes (DEGs) ranked by adjusted p-value (padj) identified in PS2N141I vs. PS2+/- mouse hippocampi are reported in Figure 4A. As an internal validation of the RNA-sequencing experiment, *Psen2* was the most significantly downregulated gene in PS2-/- relative to WT mice (log_2_FC = -1.48, padj = 2.26 x 10^-14^; Figure S3A), consistent with efficient disruption of *Psen2* mRNA expression and confirming the quality and sensitivity of the dataset. Volcano plots of DEGs are reported in Figure S3B and Figure S3C.

**Figure 4.**
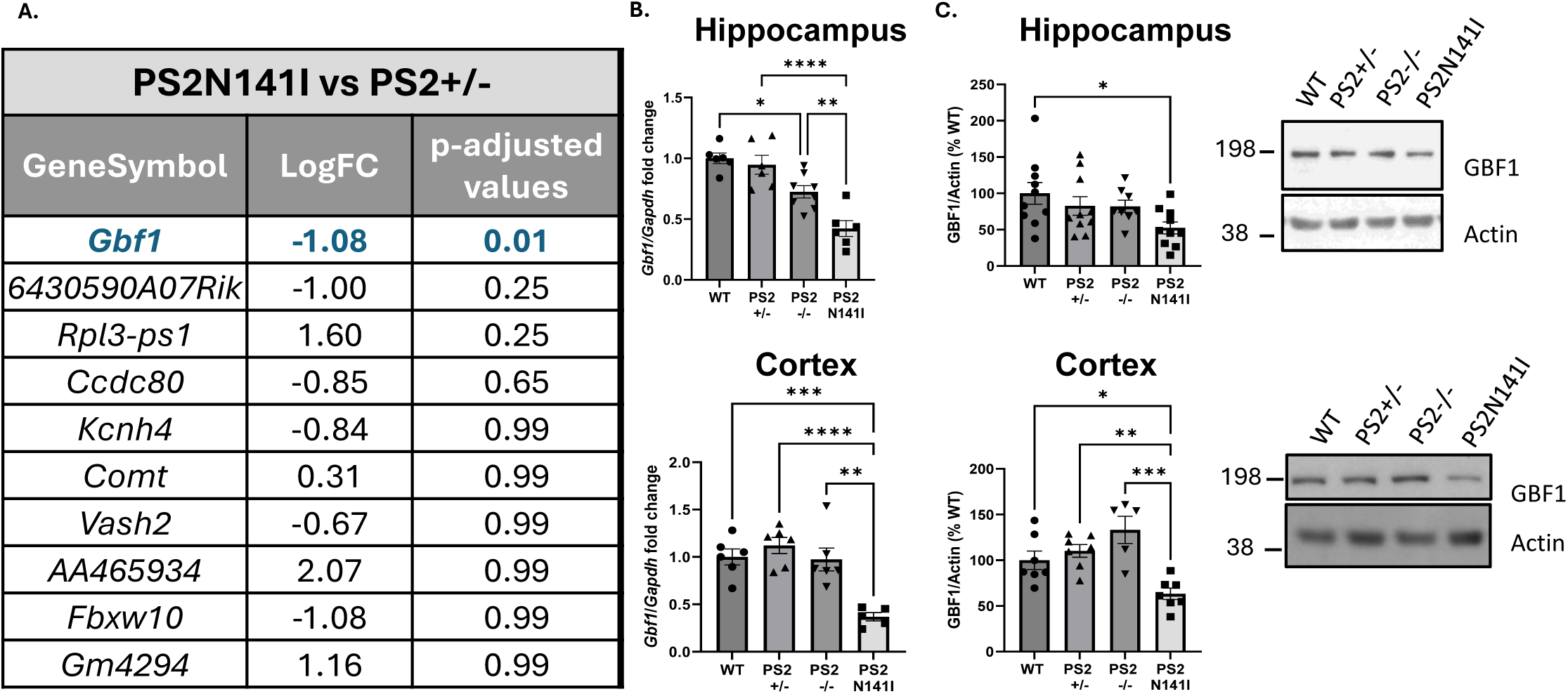
*Gbf1* is transcriptionally and post-transcriptionally downregulated in PS2N141I mouse brain. (A) Top 10 DEGs ranked by p-adjusted values in PS2N141I vs. PS2+/- mouse hippocampi at 3 months-old. (B) Measure of *Gbf1* mRNA levels in hippocampal and cortical extracts obtained from WT, PS2+/-, PS2-/-and PS2N141I mice at 3 months old. *Gapdh* was used as loading control. \**p* < 0.05, \*\**p* < 0.01, \*\*\**p* < 0.001, \*\*\*\**p* < 0.0001 (One-way ANOVA and Tukey’s multiple comparisons test, n = 6-8 mice/group). (C) Measure of GBF1 protein levels in hippocampal and cortical extracts obtained from WT, PS2+/-, PS2-/- and PS2N141I mice at 3 months old. Actin was used as loading control, and the levels in the control group (WT) were set as 100%. \**p* < 0.05, \*\**p* < 0.01, \*\*\**p* < 0.001 (One-way ANOVA and Tukey’s multiple comparisons test, n = 5-10 mice/group). Data are presented as mean ± SEM.

When PS2N141I mice were compared to PS2+/- controls, *Gbf1* (Golgi-specific Brefeldin A-resistance Guanine nucleotide Exchange Factor 1; log₂FC = -1.08, padj = 0.01) was the only gene reaching statistical significance, and notably was not among the top DEGs in the PS2-/- vs. WT comparison. GBF1 is a Golgi-resident GTP exchange factor (GEF) that activates Arf1 GTPase to drive COPI-coated vesicle formation, mediating retrograde Golgi-to-ER transport^39^. Its selective downregulation in the N141I mutation context, and not in the knockout, suggests a mutation-specific mechanism rather than a simple loss-of-function response.

*Gbf1* downregulation was independently validated by RT-qPCR in two brain regions (Figure 4B). In the hippocampus, *Gbf1* mRNA was significantly reduced in both PS2-/- and PS2N141I mice relative to WT and PS2+/-, respectively, with a significant decrease between PS2N141I and PS2-/- mice. In the cortex, the reduction was significant in PS2N141I mice compared to PS2+/-. The RT-qPCR data, obtained with greater statistical power (n = 6-8 mice/group), detected a significant *Gbf1* reduction in both PS2-/- and PS2N141I hippocampus, suggesting that the absence of a significant result for PS2-/- in the RNA-seq analysis (n = 4 mice/group) represents a false negative due to limited sample size. *Gbf1* downregulation in the hippocampus may therefore not be fully mutation-specific by contrast to the cortex.

At the protein level, GBF1 was significantly reduced in the cortex of PS2N141I mice compared with PS2+/- controls. In the hippocampus, a significant decrease was detected in PS2N141I mice relative to WT mice, but the comparison between with PS2+/- controls did not reach statistical significance (Figure 4C). This likely reflects the higher intrinsic biological variability of the hippocampus – a structurally heterogeneous region comprising distinct cellular subpopulations – which inflates inter-individual variance and reduces statistical power. Critically, GBF1 protein levels were not significantly altered in PS2-/- mice in either brain region, establishing that the protein-level effect is specific to the N141I mutation. No significant differences were observed in PS2+/- vs. WT mice, confirming that PS2 haploinsufficiency does not reduce GBF1 and the N141I effect on GBF1 levels are not due to changes in PS2 levels.

To map the anatomical source of GBF1 changes, immunohistochemical staining was performed on coronal brain sections from all four genotypes. GBF1 immunoreactivity was detected broadly across the brain and was specifically quantified in layer V of the somatosensory cortex and three hippocampal subregions: the subiculum, the dentate gyrus (DG), and Cornu Ammonis 1 (CA1). In layer V of the somatosensory cortex, the density of GBF1-positive cells (cells/mm²) was significantly reduced in PS2N141I mice compared to WT, PS2+/-, and PS2-/- mice, while PS2-/- mice did not differ significantly from WT (Figure 5A). A similar pattern was observed in the subiculum, where GBF1 cell density was significantly lower in PS2N141I mice relative to both PS2+/- and PS2-/- mice, again with no significant reduction in the PS2-/- group (Figure 5B). These two regions thus show a clear mutation-specific reduction in GBF1-expressing cell density that is not recapitulated by the knockout. In the dentate gyrus, GBF1 fluorescence intensity was significantly reduced in PS2N141I mice compared to PS2+/- mice, and in PS2-/- mice compared to WT mice (Figure 5C), mirroring the RT-qPCR findings and suggesting that both genotypes converge on reduced *Gbf1* expression in this subregion although through potentially distinct mechanisms. In the CA1, GBF1 fluorescence intensity was significantly lower in PS2N141I mice compared to both WT and PS2-/- mice, while only a trend was observed relative to PS2+/- controls, and PS2-/- mice did not differ from WT (Figure 5D). Although the most robust reductions in PS2N141I mice were detected against non-controls (WT and PS2-/-) rather than the direct PS2+/- control, likely reflecting inter-individual variability within the PS2+/- group rather than a genuine absence of effect, this data nonetheless further supports the mutation-specificity of *Gbf1* downregulation in this subregion. Taken together, the immunohistochemical data confirm and spatially resolve the *Gbf1* downregulation observed at the mRNA and protein levels, and consistently identify the N141I mutation – rather than PS2 loss-of-function – as the primary driver of GBF1 reduction across cortical and hippocampal regions. Co-immunostaining with the neuronal marker NeuN confirmed the neuronal identity of GBF1-positive cells in both the subiculum (Figure 6A) and the somatosensory cortex layer V (Figure 6B). To our knowledge, this represents the first characterization of *Gbf1* expression in the mouse brain at the regional and cell-type level.

**Figure 5.**
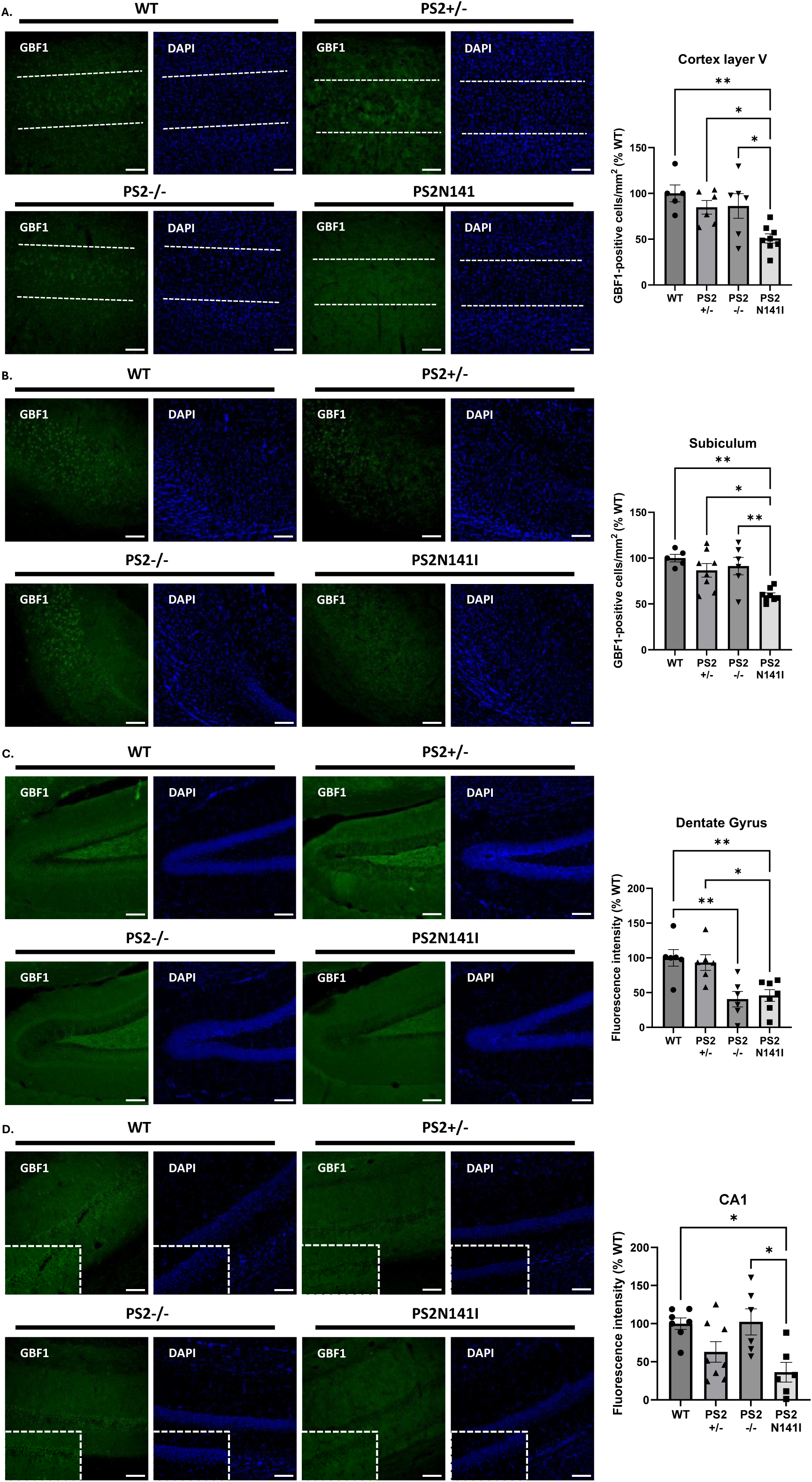
GBF1 expression is regionally reduced in PS2N141I mouse brain. (A) GBF1-positive cells were counted in layer V of the somato-sensory cortex (delimited by a white dotted line) from WT, PS2+/-, PS2-/- and PS2N141I mice at 3 months old and normalized to the area in mm^2^. The levels in the control group (WT) were set as 100%. \**p* < 0.05, \*\**p* < 0.01 (One-way ANOVA and Tukey’s multiple comparisons test, n = 5-8 mice/group). (B) GBF1-positive cells were counted in the subiculum from WT, PS2+/-, PS2-/- and PS2N141I mice at 3 months old and normalized to the area in mm^2^. The levels in the control group (WT) were set as 100%. \**p* < 0.05, \*\**p* < 0.01 (One-way ANOVA and Tukey’s multiple comparisons test, n = 6-8 mice/group). (C) GBF1 Fluorescence intensity was quantified in the dentate gyrus from WT, PS2+/-, PS2-/- and PS2N141I mice at 3 months old. The levels in the control group (WT) were set as 100%. \**p* < 0.05, \*\**p* < 0.01 (One-way ANOVA and Tukey’s multiple comparisons test, n = 6-8 mice/group). (D) GBF1 Fluorescence intensity was quantified in the CA1 from WT, PS2+/-, PS2-/- and PS2N141I mice at 3 months old. The levels in the control group (WT) were set as 100%. \**p* < 0.05 (One-way ANOVA and Tukey’s multiple comparisons test, n = 5-9 mice/group). Scale bar: 100 µm. Data are presented as mean ± SEM.

**Figure 6.**
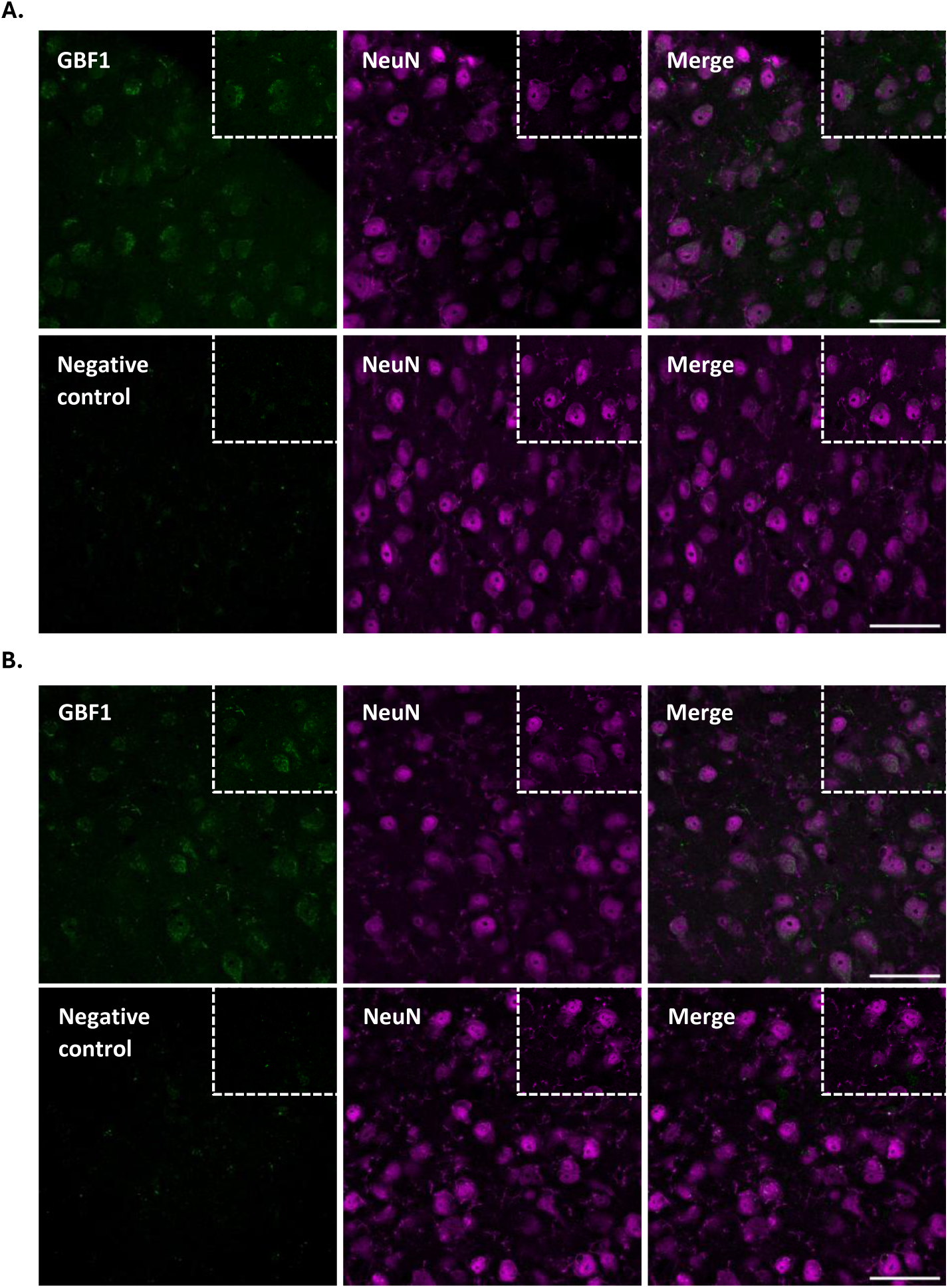
GBF1 is localized in neurons of the subiculum and somatosensory cortex layer V in the mouse brain. (A) GBF1 and NeuN co-immunostaining in the subiculum of WT mice at 3 months old. Inlets show zoomed in images. (B) GBF1 and NeuN co-immunostaining in the layer V of the somato-sensory cortex of WT mice at 3 months old. Inlets show zoomed in images. Scale bar: 50 µm.

To determine whether the *Gbf1* deficit observed *in vivo* was recapitulated in the primary neuronal model used for lipid studies, GBF1 protein levels were measured by Western blotting in primary neurons. In PS2-/- neurons, GBF1 protein levels were not significantly different from WT neurons (Figure 7A). In PS2N141I neurons, GBF1 protein levels were significantly reduced compared to PS2+/- neurons (Figure 7B). These results confirm that *Gbf1* downregulation in primary neurons is mutation specific.

**Figure 7.**
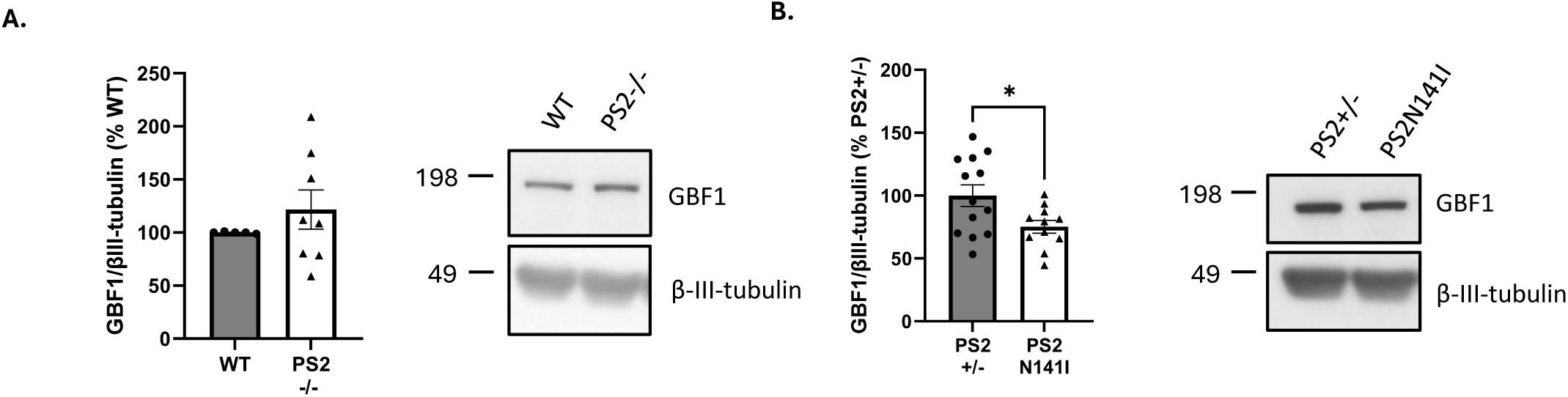
GBF1 protein levels are reduced in PS2N141I primary neurons. (A) Measure of GBF1 protein levels in WT and PS2-/- primary neurons. β-III-tubulin was used as loading control, and the levels in the control group (WT) were set as 100%. Non-significant (Unpaired t-test, N = 3 experiments, n = 5-8 embryos/group). (B) Measure of GBF1 protein levels in PS2+/- and PS2N141I primary neurons. β-III-tubulin was used as loading control, and the levels in the control group (PS2+/-) were set as 100%. \**p* < 0.05 (Unpaired t-test, N = 5 experiments, n = 11-13 embryos/group). Data are presented as mean ± SEM.

To assess whether GBF1-associated effectors were also transcriptionally dysregulated, mRNA levels of lipid droplets-associated enzymes recruited by GBF1, including *Pnpla2* (encoding ATGL; Figure S4A) and *Plin2* (encoding Perilipin 2; Figure S4B), *Arf1* (Figure S4C), and *CopI* (Figure S4D), were measured by RT-qPCR in hippocampal extracts from all four genotypes. None of these genes showed statistically significant differences across genotypes, suggesting that the transcriptional impact of the N141I mutation is selectively directed at *Gbf1* itself rather than its downstream effectors. However, as protein levels for these effectors were not assessed, their potential post-transcriptional dysregulation cannot be excluded.

Taken together, these data identify *Gbf1* as a PS2-dependent downstream target whose downregulation is specific to the N141I FAD mutation, is present at both mRNA and protein levels, and is detectable in the brain and in primary neurons. The consistent dissociation between the N141I mutation (*Gbf1* reduced) and PS2 knockout (*Gbf1* unchanged) further supports a toxic gain-of-function model and implies that the lipid deficit in PS2-/- neurons involves GBF1-independent pathways.

To directly test whether *Gbf1* downregulation is sufficient to impair lipid homeostasis, we attempted siRNA-mediated *Gbf1* knockdown in primary neurons. However, primary neurons did not tolerate the transfection conditions required for efficient siRNA delivery and *Gbf1* knockdown, resulting in substantial cell loss that precluded reliable downstream analysis. We therefore turned to wild-type (WT) mouse embryonic fibroblasts (MEFs) as an alternative cellular model in which siRNA transfection is highly efficient and well tolerated, and which share with neurons a PS2-dependent regulation of lipid metabolism^36,40^. Transfection of WT MEFs with *Gbf1*-targeting siRNAs achieved approximately 50% reduction in GBF1 protein levels relative to a non-targeting scramble control, as confirmed by Western blotting (Figure 8A). This partial knockdown is comparable in magnitude to the reduction of GBF1 observed in PS2N141I neurons (∼25%; Figure 7B) and brain tissue (hippocampus: ∼30%; cortex: ∼50%; Figure 4), making it a relevant model for functional interrogation of GBF1 loss.

**Figure 8.**
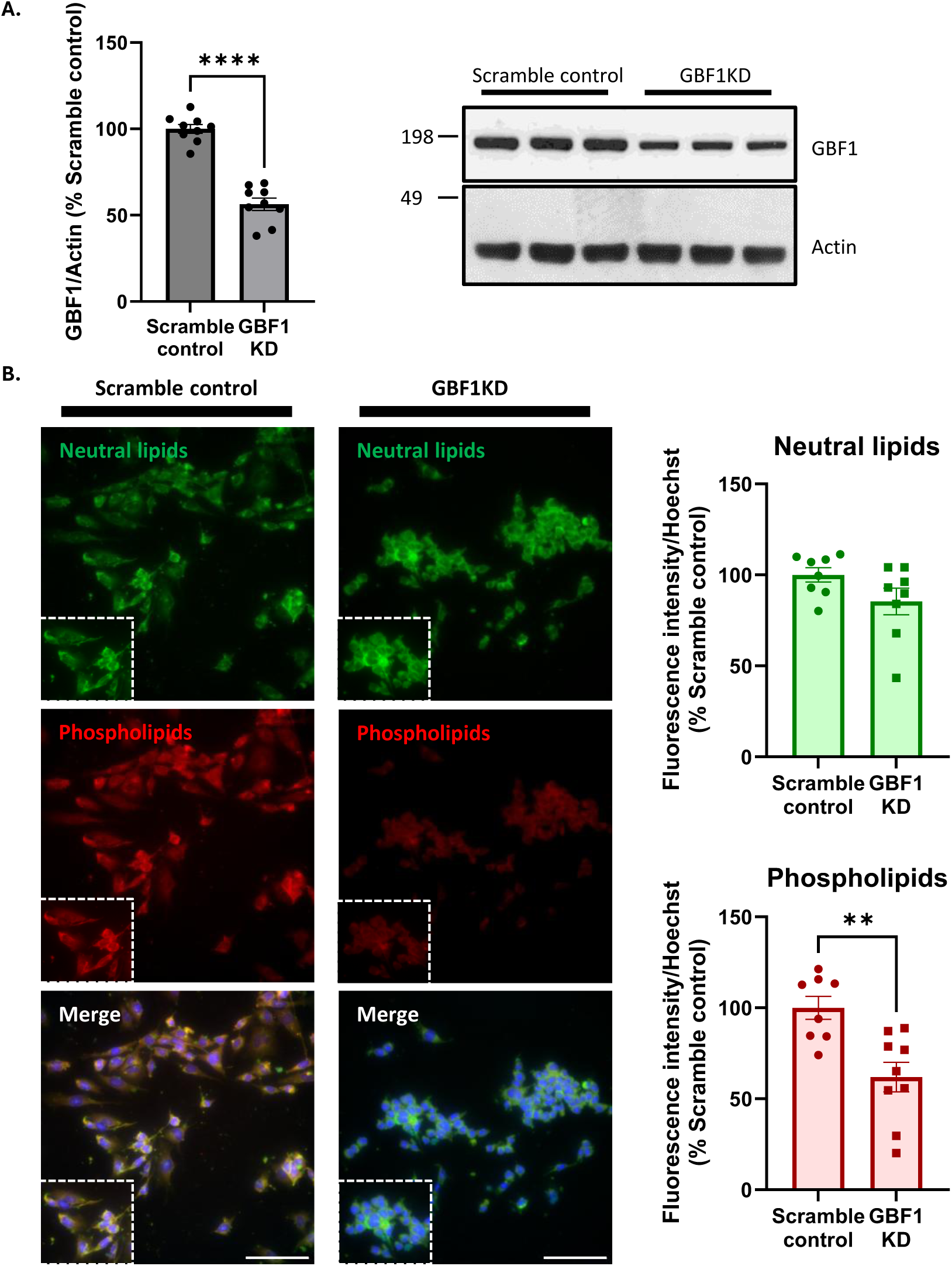
*Gbf1* knockdown selectively impairs phospholipid homeostasis in MEFs. (A) Measure of GBF1 protein levels in scramble control and GBF1KD MEFs. Actin was used as loading control, and the levels in the control group (Scramble control) were set as 100%. \*\*\*\**p* < 0.0001 (Unpaired t-test, N = 3 experiments, n = 3 wells/experiment). (B) PhenoVue Nile Red staining of scramble control and GBF1KD MEFs. Inlets show zoomed-in images. Relative fluorescence intensity (RFU) for Neutral lipids (in green) or Phospholipids (in red) was calculated and normalized to Hoechst RFU. The levels in the control group (Scramble control) were set as 100%. \*\**p* < 0.01 (Unpaired t-test, N = 3 experiments, n = 3 wells/experiment). Scale bar: 100 µm. Data are presented as mean ± SEM.

Lipid content was assessed by PhenoVue™ Nile Red staining in *Gbf1* knockdown (GBF1KD) and scramble-control MEFs. GBF1KD MEFs displayed a significant reduction in phospholipid content, whereas neutral lipid content was unaffected (Figure 8B). This selective phospholipid deficit, with no change in neutral lipids, closely mirrors the lipid profile observed in PS2N141I neurons (Figure 1B) and in LV-PSEN2N141I expressing PS2+/- primary neurons (Figure 1C). These data provide causal evidence that *Gbf1* downregulation alone is sufficient to impair phospholipid homeostasis, and support GBF1 as a functional mediator of the lipid alterations observed in the PS2N141I mutation context.

## DISCUSSION

Presenilin 2 (PS2) has long been recognized as more than the catalytic engine of the γ-secretase complex. Yet its non-amyloidogenic functions – particularly in lipid metabolism and mitochondrial homeostasis – remain far less characterized than those of PS1, and their mechanistic role to familial Alzheimer’s disease (FAD) pathogenesis is poorly understood. The present study addresses this gap by systematically comparing the consequences of complete PS2 loss and the N141I FAD mutation in primary neurons, using lipid content, transcriptomic profiling, and mitochondrial dynamics protein levels as readouts. Three convergent findings emerge. First, both the PS2 knockout and the N141I mutation reduce neuronal lipid content under basal, lipid-deprived conditions, but the lipid deficit caused by the mutation, and not by the deletion, is rescued by exogenous lipid supplementation. Second, OPA1, a key regulator of mitochondrial inner membrane fusion and cristae architecture, is specifically reduced in PS2N141I neurons under lipid-deprived conditions and is restored upon lipid supplementation. Third, the Golgi-resident guanine nucleotide exchange factor GBF1 is selectively downregulated in the N141I context at both the transcript and protein levels, in the brain and in primary neurons, while being largely unaffected by PS2 deletion. Additionally, *Gbf1* knockdown is sufficient to mimic a PS2-dependent lipid profile. Together, these results are consistent with a mutation-selective role of PS2 N141I that cannot be explained by a simple loss of function and may involve a toxic gain-of-function component that impairs lipid homeostasis through a distinct molecular mechanism, involving GBF1 and mitochondrial dynamics, that is separable from the effects of complete PS2 absence.

The divergence between PS2-/- and PS2N141I neurons upon lipid concentrate supplementation is a key observation in this study. Prior work established that PS2 FAD mutations, including N141I, produce a partial loss-of-function with respect to γ-secretase activity: they reduce γ-secretase cleavage efficiency and strongly increase the Aβ42/Aβ40 ratio, effects replicated by pharmacological γ-secretase inhibition^4,5^. The present lipid rescue data extend this partial loss-of-function concept beyond amyloidogenic processing and into lipid metabolism. We propose that in PS2-/- neurons, the complete absence of PS2 creates a metabolic deficit severe enough to prevent any response to exogenous lipid supply, characterized by a persisting phospholipid content decrease, which could be related to the decreased sphingomyelin, ceramide, and PI levels observed in PS2-/- MEFs (Figure S1). This could reflect multiple non-exclusive mechanisms: PS2 may itself participate in lipid production and lipid uptake or endocytosis, such that its total absence abrogates both endogenous lipid production and exogenous lipid internalization; alternatively, the complete loss of PS2-dependent functions at the MAM, including regulation of ER-mitochondria calcium cross-talk^22,23^, may induce a level of organellar stress incompatible with adaptive lipid responses. In contrast, PS2 N141I retains partial functionality – the mutant protein is still present and still engages its interaction partners, albeit with reduced efficiency – and this residual activity may be sufficient to permit lipid uptake and processing once substrate availability is restored. This interpretation is supported by the observation that both the lipid phenotype and the OPA1 deficit are reversible in PS2N141I neurons. Distinguishing between these possibilities will require direct measurement of lipid uptake kinetics, for example using fluorescently tagged lipid probes or FACS-based internalization assays.

The identification of *Gbf1* as a potential transcriptional target selectively downregulated by the N141I mutation represents a novel finding that provides a mechanistic link to the described phenotypes. GBF1 is a large multidomain Arf-GEF (guanine nucleotide exchange factor for ADP-ribosylation factor 1) that activates Arf1 at the cis-Golgi, thereby nucleating COPI-coated vesicle formation and maintaining the structure and function of the Golgi apparatus^41,42^. Beyond this canonical role, GBF1 and Arf1 have been implicated in a broader range of cellular functions, including lipid metabolism through GBF1’s localization at lipid droplets’ (LDs) membranes via an amphipathic helix within its HDS1 lipid-binding domain^43^. GBF1 and Arf1 regulate LD biogenesis and metabolism by controlling the recruitment of LD-associated enzymes such as the adipose triglyceride lipase ATGL (encoded by *Pnpla2*) and Perilipin 2 (PLIN2) to the LD surface^43–45^, and Arf1 has recently been shown to coordinate fatty acid metabolism and mitochondrial homeostasis through a COPI-independent pathway^46^. Beyond lipid metabolism, *Gbf1* silencing by siRNA disrupts mitochondrial morphology and membrane potential^47,48^, and GBF1-Arf1 activity has been linked to retrograde mitochondrial transport along microtubules^49^. Notably, GBF1 also participates in clathrin-independent endocytosis at the plasma membrane^50–53^, and its depletion triggers ER stress and autophagy defects^54,55^. This confluence of functions – lipid droplet regulation, mitochondrial homeostasis, and vesicular trafficking – places GBF1 at the intersection of the two phenotypic axes (lipid homeostasis, mitochondrial dynamics) observed in the present study, making it a potential mechanistic mediator of the PS2N141I phenotype. Importantly, this work demonstrates that *Gbf1* knockdown in MEF cells via siRNA mimics a lipid profile similar to that observed in PS2N141I neurons, where *Gbf1* expression was found to be downregulated. A plausible mechanistic pathway from reduced GBF1 to the observed phospholipid-selective deficit could be as follows: GBF1 loss impairs Arf1-dependent ATGL recruitment to lipid droplet surfaces, reducing lipolysis and thereby limiting fatty acid release. These fatty acids normally serve as substrates for phospholipid synthesis via the Kennedy pathway^56^. This would explain why both GBF1 knockdown in MEFs and the N141I mutation produce a phospholipid-selective deficit with neutral lipids largely unaffected, consistent with impaired fatty acid flux into phospholipid synthesis rather than a general blow in lipid metabolism. Additionally, this work provides the first characterization of *Gbf1* expression in the mouse brain, demonstrating immunoreactivity in layer V of the somatosensory cortex and in hippocampal subregions; the subiculum, the dentate gyrus, and the CA1. These regions regulate sensorimotor integration and memory consolidation and are implicated in early AD pathology. The co-localization of GBF1 with the neuronal marker NeuN confirms its neuronal expression, supporting its relevance to the lipid and mitochondrial phenotypes observed in PS2N141I primary neurons.

The mechanism by which PS2 N141I selectively reduces *Gbf1* expression – while PS2 deletion does not – is not yet established by the present data, but several plausible hypotheses can be formulated. One possibility is that the N141I mutant protein exerts a dominant gain-of-abnormal-function at the level of transcriptional regulation. PS2 is present in the nuclear envelope^57^ and has been linked to Notch and Wnt signaling cascades that influence gene expression^58^; a mutant-specific perturbation of one of these pathways could explain why the N141I substitution leads to *Gbf1* transcriptional downregulation. A second possibility involves the restricted subcellular localization of PS2: PS2-dependent γ-secretase complexes reside in late endosomes and lysosomes^28^, and the N141I mutation has been associated with endolysosomal dysfunction and impaired vesicular trafficking^27^. GBF1 itself participates in endolysosomal routing through clathrin-independent endocytosis^50–53^, and it is conceivable that a feedback loop exists whereby PS2N141I-driven endolysosomal dysfunction leads to transcriptional repression of *Gbf1* as part of a broader adaptive or maladaptive response. A third, not mutually exclusive scenario is that the N141I mutation alters calcium dynamics at the MAM in a way that is quantitatively distinct from PS2 deletion. PS2 N141I has been shown to enhance ER-mitochondria calcium shuttling, whereas PS2 loss reduces it^23,32^, potentially resulting in an altered calcium signal that differentially modulates transcriptional programs governing Golgi function. Resolving these models will require promoter activity assays, chromatin immunoprecipitation, and epistasis experiments involving manipulation of PS2 N141I.

The absence of significant mRNA changes for GBF1-associated effectors – *Arf1*, *Pnpla2*, *Plin2*, and *CopI* – in hippocampal extracts from PS2N141I mice is informative but should be interpreted cautiously. In the hippocampus at 3 months of age, the transcriptional impact of the N141I mutation appears restricted to *Gbf1* itself, suggesting that the downstream consequences of *Gbf1* downregulation are mediated through post-transcriptional or post-translational mechanisms rather than coordinated transcriptional repression of the entire Arf1-COPI pathway. This is consistent with *Arf1* being regulated primarily through GEF-mediated activation rather than transcriptional control^59,60^. However, because protein levels and activity of these effectors were not assessed, a reduction in Arf1-GTP loading, COPI recruitment efficiency, or ATGL and Perilipin 2 protein abundance at lipid droplets cannot be excluded. Protein-level and activity-level measurements of Arf1 and COPI pathway components will be necessary to fully evaluate whether *Gbf1* downregulation in PS2N141I neurons translates into functional impairment of Golgi-to-ER retrograde trafficking and lipid droplet metabolism. Moreover, characterization of GBF1-associated effectors in the cortex, where GBF1 protein reductions are most robust, or at other ages, remains to be established.

The selective reduction of OPA1 in PS2N141I neurons under lipid-deprived conditions, and its restoration upon lipid supplementation, adds a mitochondrial dimension to the PS2N141I phenotype that is mechanistically informative on multiple levels. OPA1 is the master regulator of IMM fusion and cristae architecture; its loss causes fragmentation of the mitochondrial network, loss of cristae morphology, and impairment of respiratory chain supercomplex assembly and OXPHOS efficiency^38,61,62^. Critically, OPA1 function is intimately dependent on the lipid composition of the IMM, and in particular on the mitochondria-specific phospholipid cardiolipin. Cardiolipin interacts directly with OPA1’s paddle domain through conserved motifs, stabilizing OPA1 oligomers on the membrane, and driving membrane curvature during fusion^63,64^. Recent lipidomic analysis of OPA1-deficient mouse embryonic fibroblasts confirmed that loss of OPA1 remodels the mitochondrial phospholipid profile, with shifts in phosphatidylcholine, phosphatidylethanolamine, and cardiolipin species^65^, illustrating a bidirectional relationship between lipid composition and OPA1-dependent mitochondrial morphology. In this context, the phospholipid deficit we observe in PS2N141I neurons may directly compromise OPA1 function by altering the lipid environment in which it operates, and restoration of phospholipids by exogenous lipid supplementation may re-establish the cardiolipin-enriched IMM context required for OPA1 oligomerization and membrane-remodeling activity. This is consistent with the parallel rescue of both Nile Red phospholipid signal and OPA1 protein levels upon lipid supplementation and would position the neuronal lipid deficit as upstream of – and causally contributing to – the mitochondrial phenotype. It should be noted, however, that the Nile Red phospholipid channel detects bulk membrane phospholipid fluorescence and does not resolve cardiolipin specifically; whether the observed phospholipid deficit reflects a reduction in cardiolipin cannot be determined from the present data and would require targeted cardiolipin quantification by mass spectrometry.

A unifying model emerges from these observations. We propose that PS2 N141I, acting through a combination of partial loss-of-function (evidenced by lipid rescue) and a mutation-specific gain-of-abnormal-function for *Gbf1* regulation, reduces *Gbf1* expression through a mutation-specific mechanism that does not operate in the context of complete PS2 absence. Reduced GBF1 levels impair Arf1-dependent recruitment of lipid-metabolizing enzymes to lipid droplets, thereby disrupting the regulated lipolysis and intracellular lipid trafficking that supply fatty acid substrates for phospholipid synthesis and mitochondrial membrane maintenance. The resulting phospholipid deficit compromises the lipid environment of the IMM – specifically the cardiolipin composition required for OPA1 function – potentially leading to reduced OPA1 levels or activity, impaired cristae formation, and susceptibility to mitochondrial dysfunction. This cascade is set in the background of already-established PS2-dependent deficits in MAM integrity and ER-mitochondria calcium exchange^22,23^, which would further sensitize mitochondria to perturbations in membrane lipid composition. Importantly, the present study demonstrates that *Gbf1* knockdown in MEF cells is sufficient to partially recapitulate the PS2N141I-associated lipid profile, providing proof-of-concept support for this pathway’s lipid-step in a non-neuronal context. Whether *Gbf1* knockdown in wild-type neurons similarly phenocopies the full PS2N141I phenotype, including reduced phospholipid content and OPA1 levels, remains to be established, and we identify this as a high priority for future validation of the proposed pathway.

The present findings place PS2 in the context of lipid-mitochondria coupling in neurodegeneration, a topic that has gained considerable attention in recent years. Lipid droplets in the nervous system are no longer viewed purely as inert energy stores but as dynamic organelles with protective and signaling roles under stress conditions^66^, and their dysfunction has been shown to contribute to neurodegeneration in part through defective lipid transfer to mitochondria and impaired β-oxidation^67,68^. In the AD brain, lipidomic studies have consistently documented reductions in phospholipids and sulfatides alongside increases in cholesterol esters and triglycerides, with these changes detectable in early disease stages and correlating with amyloid and tau burden^69,70^. MAMs, where PS2 is enriched, serve as hubs for precisely this type of lipid biosynthesis and transfer, synthesizing phosphatidylserine and phosphatidylethanolamine, transferring them to the mitochondria for cardiolipin remodeling, and coupling these processes to calcium-regulated Krebs cycle activity^21,71^. PS2’s dual enrichment in MAMs and late endosomes thus positions it as a regulator of lipid flow across multiple cellular compartments simultaneously, which the N141I mutation may disrupt. The observation that mitochondrial cristae morphology is altered in PS2-deficient MEF cells and that OXPHOS capacity is specifically impaired by PS2 loss – but not PS1 – in the same model^36^ suggests that the mitochondrial consequences of PS2 dysfunction can manifest independently of Aβ accumulation, supporting an Aβ-independent contribution to AD neurodegeneration.

Several important limitations of the present study merit explicit acknowledgement. First, while *Gbf1* knockdown in MEF cells recapitulated a PS2N141I-like lipid profile, thereby supporting a causal link between *Gbf1* downregulation and PS2-dependent lipid dysregulation, its involvement in the mitochondrial phenotype remains to be established. OPA1 measurement and mitochondrial morphology analysis following *Gbf1* knockdown will therefore be necessary to establish directionality within the proposed pathway. siRNA-mediated *Gbf1* knockdown in wild-type primary neurons was not achievable in the present study, likely due to the high sensitivity of primary neurons to transfection protocols and the essential cellular functions of GBF1. Ideally, neuronal *Gbf1* knockdown combined with Nile Red staining would confirm whether the MEF findings translate to a neuronal context. Second, while the Nile Red assay provides a useful global readout of neutral lipid and phospholipid content, it does not resolve individual lipid species; targeted lipidomics, with particular attention to phosphatidylserine, phosphatidylethanolamine, and cardiolipin species, would be required to determine the specific lipid changes underlying the phenotypes observed and to link them mechanistically to OPA1 function. Third, further experiments investigating mitochondrial function are necessary in addition to our data to draw conclusions about a coordinated dysregulation of the mitochondrial dynamics’ machinery. Fourth, protein level evaluation of GBF1 effectors is necessary to better understand the exact implication of the Arf1-COPI pathway. Finally, all neuronal experiments were conducted in embryonic primary cultures, which do not fully recapitulate the synaptic connections and metabolic maturity present in the adult brain; validation in adult neurons, organotypic slices, or patient-derived iPSCs would substantially extend the translational relevance of these findings.

In conclusion, this study establishes that the PS2 N141I FAD mutation exerts a partial and selective impairment of neuronal lipid homeostasis that is mechanistically distinct from the effects of complete PS2 loss, reveals a lipid-sensitive mitochondrial phenotype centered on OPA1 that is specific to the N141I mutation context, and identifies *Gbf1* as a novel PS2N141I-dependent transcriptional target expressed in neurons of the cortex and hippocampus. These findings suggest a PS2-GBF1-lipid-mitochondria axis as a potential Aβ-independent contributor to neuronal vulnerability in PS2-linked FAD, and open new avenues for investigating whether GBF1 or downstream components of this pathway could serve as therapeutic targets or early biomarkers in familial, and potentially sporadic forms, of AD.

## Supporting information

Supplemental document S1 with Figures S1-S4 and Tables S1-S4

## RESOURCE AVAILABILITY

### Lead contact

- Requests for further information and resources should be directed to and will be fulfilled by the lead contact, Pascal Kienlen-Campard.

### Materials availability

- This study did not generate new unique reagents.

### Data and code availability

- *RNA-seq data have been deposited in the NCBI Gene Expression Omnibus (GEO) database (GSE accession code: GSE328769) and are publicly available as of the date of publication*.
- *This paper does not report original code*.
- *Any additional information required to reanalyze the data reported in this paper is available from the lead contact upon request*.

## ACKNOWLEDGMENTS

We thank Esther Paître from the Aging and Dementia lab, Jean-François Geuens, and Caroline Bouzin from the 2IP Imaging Platform (Institut de Recherche Expérimentale et Clinique, IREC, UCLouvain, Brussels, Belgium) for their technical support. We also thank the team of Suman Jayadev (University of Washington, Seattle, WA, USA) for kindly providing the transgenic PS2N141I mouse model. S.Sa. was supported by an Aspirant fellowship from the F.R.S.-FNRS. N.S. was funded by a Chargé de Recherche postdoctoral fellowship from the F.R.S.-FNRS. This work was supported by grants from the SAO-FRA Alzheimer Research Foundation (SAO-FRA 2018/0025), UCLouvain Action de Recherche Concertée (ARC21/26–114), Fondation Louvain, Queen Elisabeth Medical Foundation (FMRE AlzHex), F.R.S.-FNRS (FNRS J.0106.22) attributed to P.K.C.

## AUTHOR CONTRIBUTIONS

Conceptualization, S.Sa. and P.K.C.; methodology, S.Sa., P.K.C.; Investigation, S.Sa., C.W., A.L., S.St., J.M., N.S.; writing—original draft, S.Sa.; writing—review & editing, S.Sa., C.W., A.L., S.St., J.M., G.M., N.S., P.K.C.; funding acquisition, P.K.C.; resources, P.K.C.; supervision, P.K.C.

## DECLARATION OF GENERATIVE AI AND AI-ASSISTED TECHNOLOGIES

During the preparation of this work, the author(s) used Claude Opus 4 in order to identify potential limitations and strengths of the present study. After using this tool or service, the author(s) reviewed and edited the content as needed and take(s) full responsibility for the content of the publication.

## SUPPLEMENTAL INFORMATION

**Document S1. Figures S1–S4, Tables S1–S4**

## STAR⍰METHODS

### KEY RESOURCES TABLE

| REAGENT or RESOURCE | SOURCE | IDENTIFIER |
| --- | --- | --- |
| **Antibodies** |  |  |
| Rabbit polyclonal anti-GBF1 | Abcam | ab86071 |
| Mouse monoclonal anti-NeuN | Millipore | MAB377 |
| Chicken polyclonal anti-MAP2 | Thermo Fisher Scientific | PA1-16751 |
| Goat polyclonal anti-rabbit conjugated with Alexa Fluor™ 488 | Thermo Fisher Scientific | A11034 |
| Goat polyclonal anti-mouse conjugated with Alexa Fluor™ 647 | Thermo Fisher Scientific | A21236 |
| Goat polyclonal anti-chicken conjugated with Alexa Fluor™ 647 | Thermo Fisher Scientific | A21449 |
| Rabbit monoclonal anti-Presenilin 2 | Cell Signaling | 9979S |
| Rabbit monoclonal anti-Presenilin 1 | Cell Signaling | 5643 |
| Rabbit monoclonal anti-OPA1 | Cell Signaling | 80471 |
| Rabbit monoclonal anti-Mitofusin 2 | Cell Signaling | 9482 |
| Rabbit monoclonal anti-DRP1 | Cell Signaling | 8570 |
| Rabbit polyclonal anti-β-III-tubulin | Sigma-Aldrich | T2200 |
| Rabbit polyclonal anti-actin | Sigma-Aldrich | A2066 |
| Goat polyclonal anti-rabbit conjugated with HRP | Cell Signaling | 7074 |
| **Bacterial and virus strains** |  |  |
| **Biological samples** |  |  |
| **Chemicals, peptides, and recombinant proteins** |  |  |
| Gibco™ Chemically Defined Lipid Concentrate | Thermo Fisher Scientific | 11905031 |
| Lipofectamine™ RNAiMAX Transfection Reagent | Thermo Fisher Scientific | 13778075 |
| **Critical commercial assays** |  |  |
| TruSeq Stranded mRNA kit | Illumina | 20020594 |
| PhenoVue™ Nile Red Lipid Stain | Revvity | CP41 |
| TriFECTa DsiRNA Duplex targeting mouse *Gbf1* | IDT | N/A |
| **Deposited data** |  |  |
| RNA-Seq data | This paper | GEO: GSE328769 |
| **Experimental models: Cell lines** |  |  |
| Mouse: Mouse Embryonic Fibroblasts (MEFs) | ATCC | CRL-2991 |
| Human: Human Embryonic Kidney 293T (HEK293T) | ATCC | CRL-3216 |
| **Experimental models: Organisms/strains** |  |  |
| Mouse: PS2-/-; B6.129P-Psen2tm1Bdes/J | The Jackson Laboratory | JAX:005617 |
| Mouse: PS2N141I; transgenic PSEN2 N141I mice | Fung *et al.*<sup>33</sup> | N/A |
| **Oligonucleotides** |  |  |
| Primers for DsiRNA transfection, see Table S1 | This paper | N/A |
| Primers for RT-qPCR, see Table S3 | This paper | N/A |
| **Recombinant DNA** |  |  |
| pCMV delta R8.2 | Addgene | 12263 |
| pMD2.G | Addgene | 12259 |
| pLenti CMV/TO Puro (w175-1) | Addgene | 17482 |
| pCMV-PSEN2WT | This paper | N/A |
| pCMV-PSEN2N141I | This paper | N/A |
| **Software and algorithms** |  |  |
| ImageJ | Schneider et al.<sup>72</sup> | https://imagej.net/ij/ |
| GraphPad Prism 10.0 | GraphPad Software, Boston, Massachusetts, USA | https://www.graphpad.com/ |
| **Other** |  |  |
| Nikon Eclipse Ni-L upright fluorescence microscope | Nikon Instruments | https://www.microscope.healthcare.nikon.com/products/upright-microscopes/eclipse-ni-series |
| Olympus Fluoview FV1000 laser scanning confocal microscope | Olympus | https://www.olympusamerica.com/ |

## EXPERIMENTAL MODEL AND STUDY PARTICIPANT DETAILS

### Animals

Two independent mouse models were used in this study. First, PS2 knockout (PS2-/-) mice (#005617)^73^ were obtained from The Jackson Laboratory (Bar Harbor, ME, USA) and compared to wild-type (WT) littermates. Second, PS2N141I transgenic mice were kindly provided by Dr. Suman Jayadev (University of Washington, Seattle, WA, USA)^33^. Briefly, the human *PSEN2* gene carrying the N141I mutation is expressed under a chicken-β-actin promoter on a PS2 heterozygous knockout (PS2+/-) background. Breeding was performed by crossing WT mice with PS2-/- mice expressing the PS2N141I transgene, yielding two littermate genotypes: PS2+/- mice without the transgene, used as controls, and PS2+/- mice carrying the PS2N141I transgene, used as the experimental group. All mouse lines were maintained on a C57BL/6J genetic background. For all experiments, age- and sex-matched littermates were used as (WT vs. PS2-/-; PS2+/- vs. PS2N141I). Sex had no significant effect on experimental outcomes, and results were obtained in both sexes. Genotyping was performed by polymerase chain reaction (PCR) on genomic DNA extracted from tail biopsies at weaning.

Animals were housed on a 12 h light/12 h dark cycle in a specific pathogen-free facility with *ad libitum* access to food and water. All experimental procedures were conducted in accordance with the regulations of the UCLouvain Institutional Animal Care and Use Committee (2021/UCL/MD/022).

### Primary Neuronal Cultures

Mouse primary cortical and hippocampal neurons were isolated from embryonic day 19 (E19) embryos as previously described^74^. Briefly, embryonic brains were rapidly dissected in ice-cold Hank’s Balanced Salt Solution (HBSS; Thermo Fisher Scientific, Waltham, MA, USA) supplemented with 0.2% D-glucose. Cortices and hippocampi were isolated under a stereomicroscope and meninges were removed. Tissues were mechanically dissociated by sequential trituration: 15 strokes with a glass pipette, followed by 10 strokes with a flame-narrowed glass pipette. After sedimentation for 5 min at room temperature, the supernatant was layered onto a bed of Fetal Bovine Serum (FBS; Thermo Scientific, Rockford, IL, USA) and centrifuged at 1,000 × *g* for 10 min. The pellet was resuspended in complete Neurobasal® (NB; Thermo Fisher Scientific) medium supplemented with 2% (v/v) B-27™ supplement, 1 mM L-glutamine, 5 mM glucose, and 1% (v/v) penicillin/streptomycin (100 U/ml and 100 g/ml, respectively). Neurons were plated at the preferred density onto pre-coated poly-L-lysine (0.1 mg/ml; Sigma-Aldrich) dishes and maintained at 37 °C in a humidified atmosphere containing 5% CO_2_.

Where indicated, neurons were supplemented at days *in vitro* 10 (DIV10) with the Gibco™ Chemically Defined Lipid Concentrate (#11905031; Thermo Fisher Scientific; final dilution 1:100 in complete Neurobasal® medium). All cultures were collected at DIV13 for downstream analyses.

### Cell Lines

Mouse embryonic fibroblasts (MEFs) wild-type (WT), PS1 knockout (PS1-/-), and PS2 knockout (PS2-/-) were cultured in DMEM medium (Life Technology Corporation, Carlsbad, CA, USA) supplemented with 10% heat-inactivated FBS and 1% penicillin/streptomycin. All cell cultures were maintained at 37 °C in a humidified atmosphere containing 5% CO_2_.

## METHOD DETAILS

### Lentiviral Production and Neuronal Infection

To express human PS2 wild-type or the N141I mutant in primary neurons independently of the transgenic background, lentiviral vectors were used. The coding sequences of human *PSEN2* wild-type and *PSEN2* N141I were subcloned into the pLenti CMV/TO Puro (w175-1) transfer vector (Addgene #17482) to generate pCMV-PSEN2WT and pCMV-PSEN2N141I, respectively.

Lentiviral particles were generated as previously described^75^. Briefly, HEK293T cells at 70-80% confluence were co-transfected with the transfer plasmid (pCMV-PSEN2WT or pCMV-PSEN2N141I), the VSV-G envelope plasmid pMD2.G (Addgene #12259), and the packaging plasmid pCMV-dR8.2 (Addgene #12263). 48 h after transfection, conditioned medium and detached cells were harvested together and centrifuged at 1,500 ×□*g* for 10 min at 4 °C to remove cellular debris. The clarified supernatant was filtered through a 0.45 µm membrane and concentrated using LentiX™ Concentrator reagent (Clontech Laboratories, Mountain View, CA, USA) according to the manufacturer’s instructions. Concentrated viral particles were resuspended in complete Neurobasal® medium and stored at – 80 °C until use.

PS2+/- primary neurons were infected at DIV6 by addition of lentiviral particles directly to the culture medium. 24 h after infection, the entire culture medium was replaced with fresh complete Neurobasal® medium. PS2 expression was confirmed at DIV13 by Western blotting.

### siRNA Transfection

Prior to siRNA transfection, MEF cell medium was changed to antibiotic-free medium. *Gbf1* knockdown was achieved using the TriFECTa DsiRNA Duplex Kit (Integrated DNA Technologies, Coralville, IA, USA) targeting mouse *Gbf1*, according to the manufacturer’s instructions. A non-targeting scramble DsiRNA duplex included in the kit was used as a negative control. Briefly, scramble control or *Gbf1*-targeting siRNAs (see Table S1) were mixed with Lipofectamine™ RNAiMAX Transfection Reagent (Thermo Fisher Scientific) at a 1:2 ratio in Gibco™ Opti-MEM™ (Thermo Fisher Scientific) to obtain a final siRNA concentration of 10 nM. The mixture was added to the cells for 24 hours, after which the medium was replaced, and the same transfection was repeated. This process was carried out every 24 hours for 3 days until collection.

### Mass Spectrometry

WT, PS1-/- and PS2-/- MEFs were seeded and 48 h after cells have been washed and scraped off in PBS and centrifuged for 5 min at 10,000 rpm. Pellets were sonicated in lysis buffer (125mM Tris pH 6.8, 20% glycerol, 4% SDS) with complete protease inhibitor cocktail. The lipid content was analyzed by HPLCl1MS after liquid/liquid extraction of the lipids in the presence of the appropriate internal standards, following established protocols^76,77^. Following fractionation of the lipid extract using solid phase extraction, the lipid content was analyzed on an LTQl1Orbitrap mass spectrometer coupled to an Accela HPLC system. Lipid separation was performed using a Cl118 Supelguard prel1column and a Kinetex LCl118 column (5 μm, 4.6 × 150 mm). Three mobilel1phase systems were used according to lipid class of interest namely (i) sphingomyelins, (ii) phospholipids, and (iii) ceramides^76^. The signal (AUC) of each lipid was normalized to the signal (AUC) of its internal standard. The data are reported relative to the WT condition set at 100%.

### Immunofluorescence

#### Mouse brain sections

Mice were anaesthetized and transcardially perfused with ice-cold phosphate-buffered saline (PBS). Brains were post-fixed in 4% paraformaldehyde (PFA) at 4 °C for 24 h, cryoprotected in PBS containing 30% sucrose with 0.02% sodium azide and stored at 4 °C. 30 µm-thick coronal brain sections were cut using a freezing microtome and collected as free-floating sections in a cryoprotectant solution (30% ethylene glycol, 20% glycerol) at – 20 °C until use.

For immunostaining, free-floating sections were rinsed three times in PBS and blocked/permeabilized for 30 min at room temperature in PBS containing 3% Bovine Serum Albumin (BSA) and 0.5% Triton X-100. Brain sections were then incubated overnight at 4 °C with primary antibodies diluted in blocking buffer (see Table S2). After three washes with PBS, sections were incubated for 1 h at room temperature with species-appropriate fluorophore-conjugated secondary antibodies and Hoechst 33342 nuclear stain (1:10,000; #62249; Thermo Fisher Scientific), all diluted in blocking buffer. Finally, sections were washed three times in PBS and mounted on Superfrost™ Plus slides, allowed to air-dry, and coverslipped with Mowiol 4-88 mounting medium.

Images were acquired using either a Nikon Eclipse Ni-L upright fluorescence microscope or an Olympus Fluoview FV1000 laser scanning confocal microscope. GBF1 immunoreactivity was quantified using ImageJ (NIH): cells displaying a clear intracellular GBF1 signal were manually counted in each region of interest (layer V of the somatosensory cortex, subiculum) and the count was normalized to the area of the region (mm^2^). For the Cornu Ammonis 1 (CA1) and the dentate gyrus (DG), the integrated density of the fluorescence signal was measured within each acquired field, local background was subtracted, and the result was expressed as relative fluorescence units (RFU).

#### Primary neurons and cell lines

Neurons and MEFs grown on glass coverslips were fixed for 10 min in 4% PFA at room temperature, washed in PBS, and stored at 4 °C in PBS containing 0.02% sodium azide until use.

For neurons’ immunostaining, cells were permeabilized with a solution of PBS/0.3% Triton X-100 for 30 min, and non-specific sites were blocked with PBS/0.3% Triton/5% FBS for 30 min. Primary antibodies diluted in the blocking solution were applied overnight at 4 °C (see Table S2). After 3 washes of 10 min with PBS, cells were incubated with secondary antibodies and Hoechst 33342 (1:10,000) diluted in blocking buffer.

Lipid staining was performed by incubating the cells (neurons and MEFs) with PhenoVue™ Nile Red Lipid Stain (1:1,000; Revvity) for 30 min at room temperature. Cells were then consecutively washed in PBS and ultrapure water and mounted on glass slides with Mowiol 4-88.

Images were acquired on a Nikon Eclipse Ni-L fluorescence microscope. Nile Red fluorescence was quantified in ImageJ; the integrated density of the fluorescence signal was measured within each acquired field, the local background was subtracted, and the resulting relative fluorescence unit (RFU) value was normalized to the corresponding Hoechst 33342 integrated density to correct for differences in cell number.

### Quantitative RT-PCR

Total RNA was extracted from mouse brain tissue using TriPure™ Isolation Reagent (Roche, Basel, Switzerland) according to the manufacturer’s instructions. RNA was resuspended in DEPC-treated water and quantified by UV absorbance at 260 nm using a BioSpec-nano spectrophotometer (Shimadzu Biotech, Kyoto, Japan); only preparations with A₂₆₀/A₂₈₀ ratios ≥ 1.8 were used.

Reverse transcription (RT) was carried out using the iScript™ cDNA Synthesis Kit (Bio-Rad Laboratories, Hercules, CA, USA) with 1 μg of total RNA in a 20 μl reaction. Quantitative PCR (qPCR) was performed for the amplification of cDNAs using gene-specific primers (Sigma-Aldrich; see Table S3) and the GoTaq® qPCR Master Mix (Promega, Madison, WI, USA) on a real-time PCR system. Relative mRNA levels were calculated using the 2^−ΔΔCT^ method, with *Gapdh* as the housekeeping reference gene. Results are expressed as fold changed normalized to the mean of the control group.

### RNA-Sequencing

#### Sample preparation

Hippocampal tissues were collected from WT, PS2+/-, PS2-/-, and PS2N141I mice at 3 months of age (n = 4 animals per genotype; 16 animals in total). Tissues were homogenized in TriPure™ Isolation Reagent (Roche) and total RNA was extracted according to the manufacturer’s protocol. RNA was further purified using the ReliaPrep_™_ RNA Tissue Miniprep System (Promega). Concentration and purity were assessed spectrophotometrically (BioSpec-nano; Shimadzu Biotech) and RNA integrity was evaluated using the Agilent RNA 6000 Nano Kit on an Agilent 2100 Bioanalyzer (Agilent Technologies, Santa Clara, CA, USA); only samples with RNA Integrity Number (RIN) ≥ 8.0 were included. Samples (200 ng/µl) were sent to Macrogen Europe B.V. (Amsterdam, Netherlands) for library preparation and paired-end sequencing.

#### Bioinformatic analysis

Raw FASTQ files were quality checked with FastQC and adapter sequences and low-quality bases were trimmed using Trimmomatic (v0.39)^78^. Trimmed reads were aligned to the mouse reference genome (GRCm38/mm10) using HISAT2 (v2.2.1)^79^. Gene-level read counts were quantified from the aligned BAM files using featureCounts (Subread v2.0.3)^80^ with the Ensembl annotation file *Mus_musculus.GRCm38.94.gtf*. Only uniquely mapping reads were retained. Differential gene expression analysis was performed with DESeq2 (Bioconductor v1.50.2)^81^ using the Wald test; p-values were adjusted for multiple comparisons using the Benjamini–Hochberg procedure. Genes were considered differentially expressed at an adjusted p-value (padj) < 0.05.

### Protein Extraction

#### Brain tissue

Dissected brain regions were homogenized by probe sonication in ice-cold lysis buffer (20 mM Tris-HCl pH 8.0, 150 mM NaCl, 1% NP-40, 10% glycerol) supplemented with protease and phosphatase inhibitor cocktails (Roche, Basel, Switzerland). Lysates were centrifuged at 16,100 x *g* for 15 min at 4 °C, and the supernatants were collected and stored at – 80 °C until use.

#### Primary neurons and cell lines

Cells were scraped in ice-cold lysis buffer (50 mM Tris-HCl pH 7.5, 150 mM NaCl, 2 mM EDTA, 1% NP-40) supplemented with protease and phosphatase inhibitor cocktails (Roche, Basel, Switzerland). Lysates were sonicated and centrifuged at 16,100 x *g* for 10 min at 4 °C. Supernatants were collected and stored at – 80 °C until use.

### Western Blotting

Protein concentration was determined using the Pierce™ BCA Protein Assay Kit (Thermo Fisher Scientific). Equal amounts of protein (15 µg) were denatured in NuPAGE™ LDS Sample Buffer (Thermo Fisher Scientific) containing 50 mM dithiothreitol (DTT) at 70 °C for 10 min.

Proteins were separated by SDS-PAGE electrophoresis on precast NuPAGE™ 4–12% Bis–Tris gels (Thermo Fisher Scientific) using MES-SDS running buffer (Thermo Fisher Scientific). SeeBlue™ Plus2 pre-stained protein ladder (Thermo Fisher Scientific) was included as a standard. Proteins were transferred onto 0.1 μm pore-sized nitrocellulose membranes (Thermo Fisher Scientific) for 2 h at 30 V in NuPAGE™ transfer buffer (Thermo Fisher Scientific).

Membranes were blocked for 30 min at room temperature in 5% non-fat dried milk dissolved in PBS/0.1% Tween® 20 (PBS-T), then incubated overnight at 4 °C with primary antibodies diluted in PBS-T (see Table S4). After three 10 min washes in PBS-T, membranes were incubated for 1 h at room temperature with species-appropriate HRP-conjugated secondary antibodies (Sigma-Aldrich) diluted in PBS-T. Immunoreactive bands were detected by enhanced chemiluminescence (ECL). Band intensities were quantified using ImageJ (NIH) and normalized to the loading control (β-III-tubulin for primary neurons; actin for brain tissue lysates and MEFs).

## QUANTIFICATION AND STATISTICAL ANALYSIS

All statistical analyses were performed using GraphPad Prism version 10.0 (GraphPad Software, San Diego, CA, USA). Data normality was assessed with the Shapiro-Wilk test prior to selecting the appropriate statistical test. For comparisons between two groups, a parametric unpaired Student’s *t*-test or its non-parametric equivalent (Mann-Whitney U test) was applied depending on whether normality was met. For comparisons involving more than two groups, one-way or two-way ANOVA followed by the indicated post hoc multiple comparisons test was used for normally distributed data; the non-parametric Kruskal–Wallis test followed by Dunn’s multiple comparisons test was applied otherwise. The specific test used for each dataset is stated in the corresponding figure legend. Significance thresholds: \**p* < 0.05; \*\**p* < 0.01; \*\*\**p* < 0.001; \*\*\*\**p* < 0.0001. Data are expressed as mean ± SEM. The number of independent biological replicates (animals or embryos) is denoted *n*; the number of independent experimental repetitions is denoted *N*, as specified in each figure legend.

