## Supplemental document S1 with Figures S1-S4 and Tables S1-S4 for "Alzheimer’s disease-associated Presenilin 2 N141I mutation impairs neuronal lipid homeostasis and mitochondrial dynamics through selective downregulation of the Golgi exchange factor *Gbf1*"

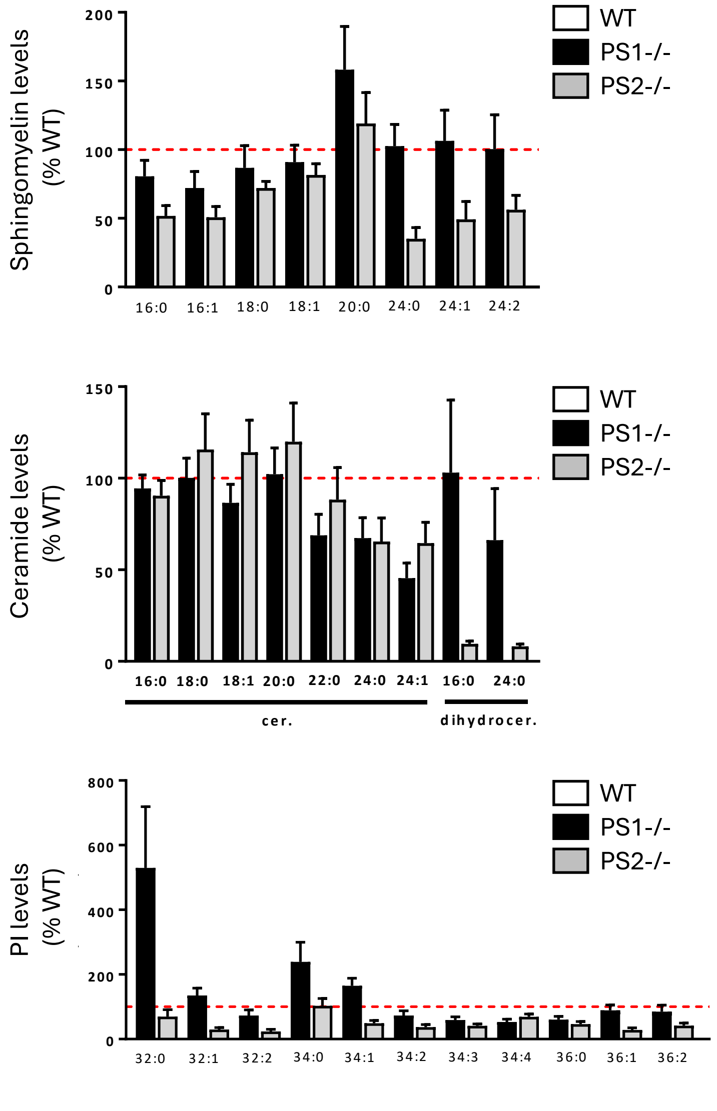


***Figure S1. S*phingomyelin, ceramide and phosphatidylinositol levels depend on PS2 in MEF cells.** Sphingomyelin, ceramides and dihydroceramides and phosphatidylinositol (PI) levels have been measured in PS1-/- and PS2-/- cells compared to those found in WT (dashed red lines). For sphingomyelin and ceramide, the first number represents the length of the acyl chain, the second number represents the number of insaturations; for PIP, the first number represents the sum of the sn1 and sn2 acyl chain length, the second number represents the total number of insaturations. The levels in the control group (WT) were set as 100%. *$p < 0.05$ (vs. WT) (One-Way ANOVA and Bonferroni's multiple comparisons test, n = 4-6).
Data are presented as mean ± SEM.

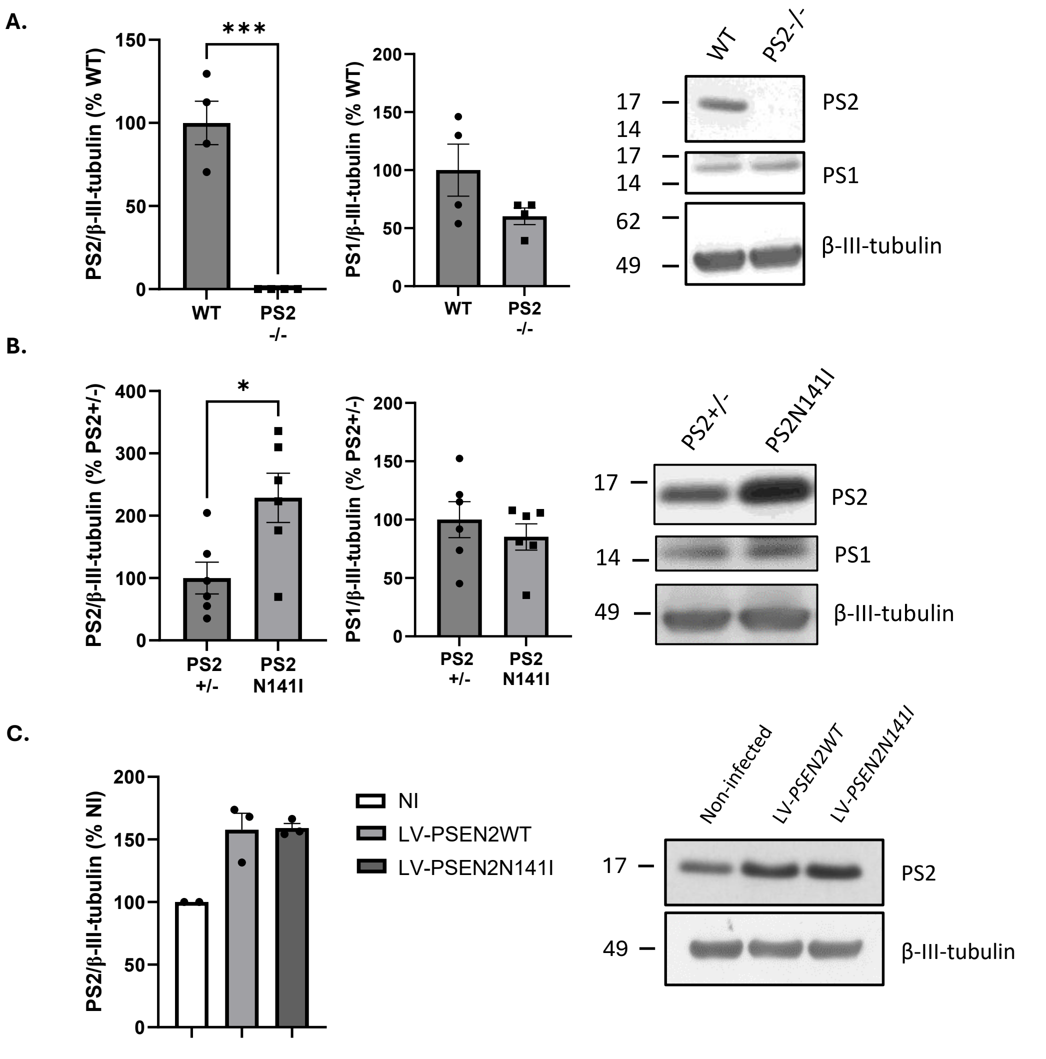


***Figure S2.*** **PS1 and PS2 levels in WT vs. PS2-/-, PS2+/- vs. PS2N141I and lentivirus-infected PS2+/- primary neurons.**

**(A)** Measure of PS1 and PS2 levels in primary neurons obtained from WT and PS2-/- E19 embryos at DIV13 (Days *In Vitro*). The levels in the control group (WT) were set as 100%. β-III-tubulin was used as loading control. ***$p < 0.0005$ (Unpaired t-test, N = 2 cultures, n = 4 embryos/group).

**(B)** Measure of PS1 and PS2 levels in primary neurons obtained from PS2+/- and PS2N141I E19 embryos at DIV13 (Days *In Vitro*). The levels in the control group (PS2+/-) were set as 100%. β-III-tubulin was used as loading control. *$p < 0.05$ (Unpaired t-test, N = 3 cultures, n = 6 embryos/group).

**(C)** Measure of PS2 levels in primary neurons obtained from WT neurons either non-infected (NI) or infected with a lentivirus (LV) expressing PS2 WT (LV-PSEN2WT) or PS2N141I (LV-PSEN2N141I) E19 embryos at DIV13 (Days *In Vitro*). The levels in the control group (NI) were set as 100%. β-III-tubulin was used as loading control. Non-significant (Unpaired t-test, N = 2 cultures, n = 2-3 embryos/group).

Data are presented as mean ± SEM.

**A.**

| PS2-/- vs WT | | |
|---|---|---|
| **GeneSymbol** | **LogFC** | **p-adjusted values** |
| *Catspere1* | 6.49 | 8.29E-26 |
| *Psen2* | -1.48 | 2.26E-14 |
| *Catspere2* | 1.89 | 3.18E-12 |
| *Slc30a10* | -0.68 | 1.20E-3 |
| *Lyplal1* | 1.06 | 2.27E-3 |
| *Ttr* | 7.54 | 9.80E-3 |
| *Kcnk2* | -0.54 | 0.02 |
| *F5* | 5.19 | 0.23 |
| *Myl1* | -1.56 | 0.29 |
| *Sccpdh* | -0.46 | 0.29 |

**B.**

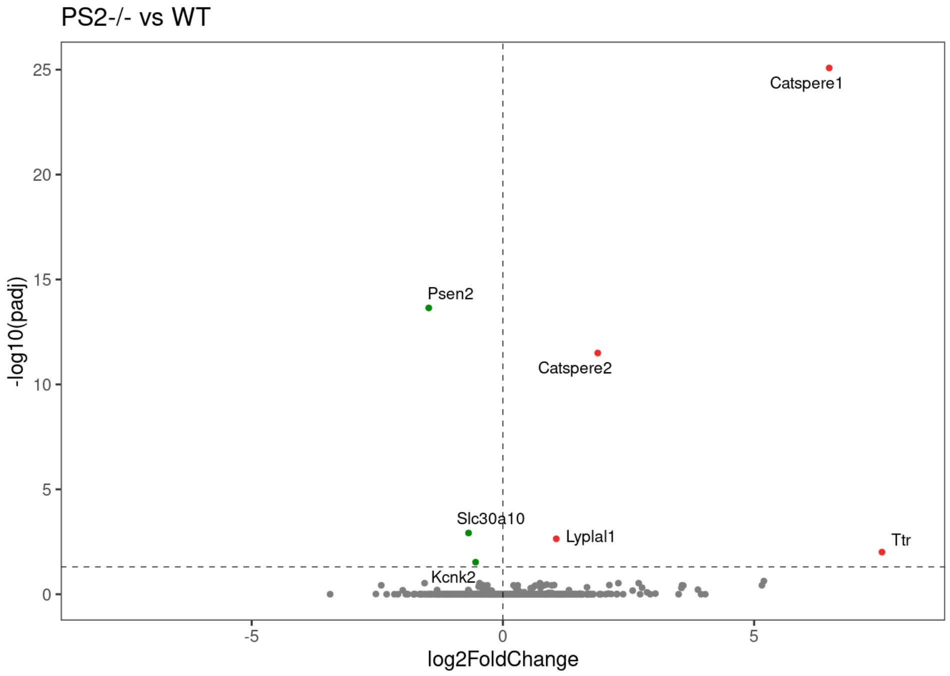


**C.**

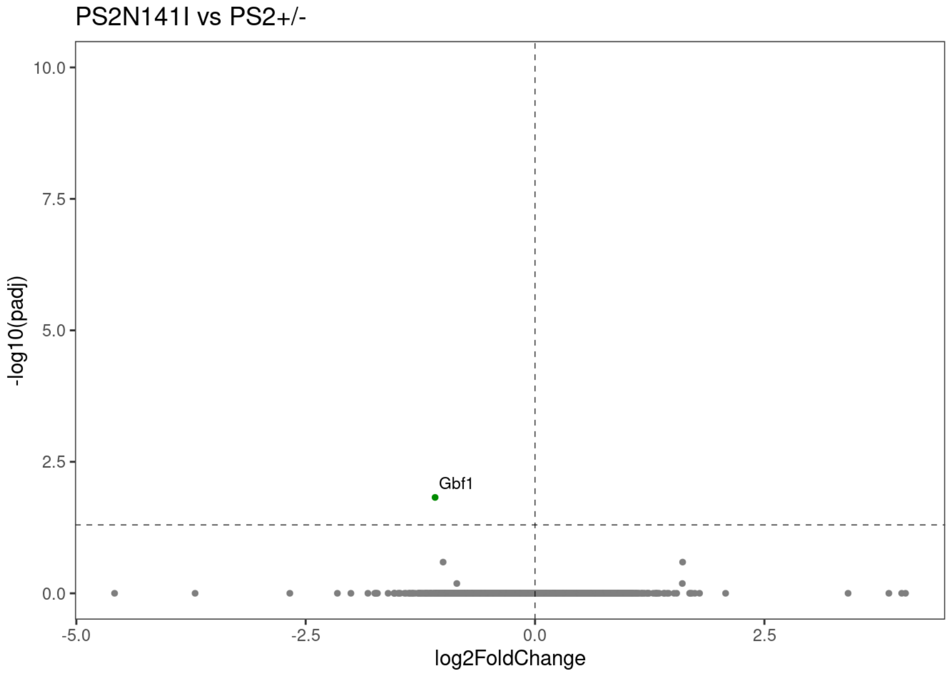


***Figure S3.*** **Differential gene expression analysis in WT vs. PS2-/- and PS2+/- vs. PS2N141I mouse hippocampus at 3 months-old.**
**(A)** Top 10 DEGs ranked by p-adjusted values in PS2-/- vs. WT mouse hippocampi at 3 months-old.
**(B)** Volcano plot comparing PS2-/- vs. WT hippocampi. The x-axis represents log2 fold change, and the y-axis shows -log10 adjusted p-value (padj). Significantly differentially expressed genes are highlighted: upregulated genes in red and downregulated genes in green. Notable genes include *Caspere1* and *Caspere2* (upregulated), and *Psen2* (downregulated). Dashed lines indicate significance and fold-change thresholds.
**(C)** Volcano plot comparing PS2N141I vs. PS2+/- hippocampi. The x-axis represents log2 fold change, and the y-axis shows −log10 adjusted p-value (padj). Significantly differentially expressed genes are highlighted with *Gbf1* identified as significantly downregulated in green. Thresholds for significance are indicated by dashed lines.

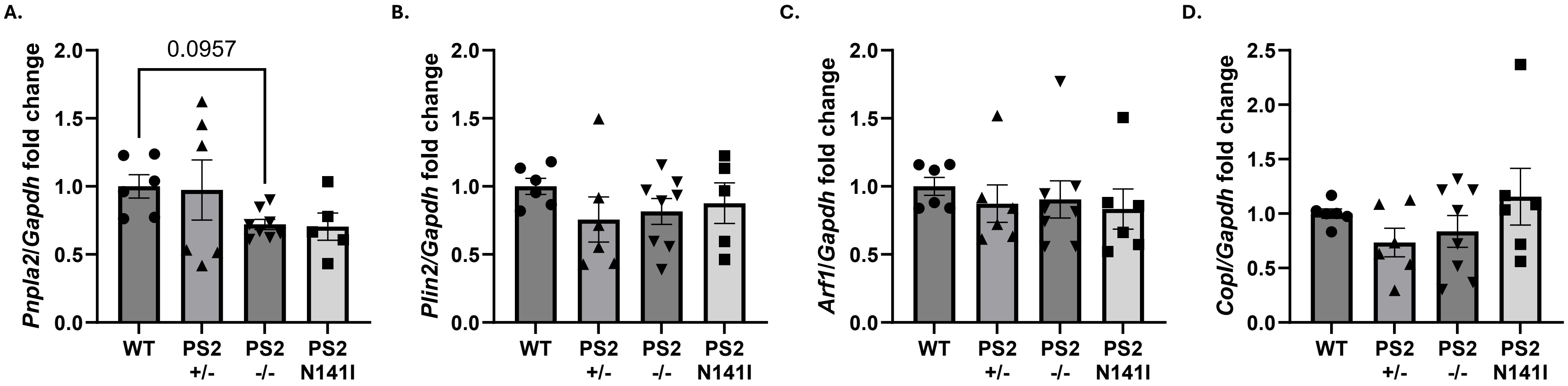


***Figure S4. Pnpla2*, *Plin2*, *Arf1* and *CopI* levels are unchanged in PS2N141I mouse hippocampus.**

**(A)** Measure of *Pnpla2* mRNA levels in hippocampal extracts obtained from WT, PS2+/-, PS2-/- and PS2N141I mice at 3 months old. *Gapdh* was used as loading control. Numeric p-value as indicated (Brown-Forsythe ANOVA and Dunnett's T3 multiple comparisons test, n = 6-8 mice/group).

**(B)** Measure of *Plin2* mRNA levels in hippocampal extracts obtained from WT, PS2+/-, PS2-/- and PS2N141I mice at 3 months old. *Gapdh* was used as loading control. Non-significant (One-way ANOVA and Tukey's multiple comparisons test, n = 6-8 mice/group).

**(C)** Measure of *Arf1* mRNA levels in hippocampal extracts obtained from WT, PS2+/-, PS2-/- and PS2N141I mice at 3 months old. *Gapdh* was used as loading control. Non-significant (Kruskal-Wallis test and Dunn's multiple comparisons test, n = 6-8 mice/group).

**(D)** Measure of *CopI* mRNA levels in hippocampal extracts obtained from WT, PS2+/-, PS2-/- and PS2N141I mice at 3 months old. *Gapdh* was used as loading control. Non-significant (One-way ANOVA and Tukey's multiple comparisons test, n = 6-8 mice/group).

Data are presented as mean ± SEM.

**Table S1. DsiRNA sequences.**

| Target | Sequence (5' – 3') | Catalogue no. | Supplier |
|---|---|---|---|
| DsiRNA duplex 1 – *Gbf1* sense | GUCAGGAUGGUGGAUAAGAA UAUTT | 512852697 | IDT |
| DsiRNA duplex 1 – *Gbf1* antisense | AAAUAUUCUUAUCCACCAUCC UGACAA | 512852697 | IDT |
| DsiRNA duplex 2 – *Gbf1* sense | GGAUGACAUUGAUAACUCCAA ACCA | 512852700 | IDT |
| DsiRNA duplex 2 – *Gbf1* antisense | UGGUUUGGAGUUAUCAAUGU CAUCCUU | 512852700 | IDT |
| DsiRNA duplex 3 – *Gbf1* sense | GGUGGACUUUUUAUGGUAUA AUAAA | 512852703 | IDT |
| DsiRNA duplex 3 – *Gbf1* antisense | UUUAUUAUACCAUAAAAAGUC CACCGU | 512852703 | IDT |
| DsiRNA negative control | Proprietary | 51-01-14-03 | IDT |

**Table S2. Primary and secondary antibodies used for immunofluorescence.**

| Target | Host species | Dilution | Catalogue no. | Supplier |
|---|---|---|---|---|
| Anti-chicken AF647 | Goat | 1:500 | A21449 | Thermo Fisher |
| Anti-mouse AF647 | Goat | 1:500 | A21236 | Thermo Fisher |
| Anti-rabbit AF488 | Goat | 1:500 | A11034 | Thermo Fisher |
| GBF1 | Rabbit | 1:200 | ab86071 | Abcam |
| MAP2 | Chicken | 1:1,000 | PA1-16751 | Thermo Fisher |
| NeuN | Mouse | 1:100 | MAB377 | Millipore |

**Table S3. Primers used for RT-qPCR.**

| Target | Forward (5' – 3') | Reverse (5' – 3') |
|---|---|---|
| Arf1 | TGGGAGCGAAACCAACG | CATGTTTGTGGACAGGTGGA |
| CopI | AAAGTCCTCCCTACCATCCC | TGTAAGTGCTTGCTGCTTGA |
| Gapdh | ACCCAGAAGACTGTGGATGG | CTGCTTCACCACCTTCTTGA |
| Gbf1 | GGCACAGTGATGAAGAGGAG | GTGTGCAGGGTATGCATCAG |
| Plin2 | AACAACACGCCTCTCAACTG | TTGGGTTCAATCATCCGCTT |
| Pnpla2 | CAACGCCACTCACATCTACG | AATGTTGGCACCTGCTTCA |

**Table S4. Primary and secondary antibodies used for Western blotting.**

| Target | Host species | Dilution | Catalogue no. | Supplier |
|---|---|---|---|---|
| β-III-tubulin | Rabbit | 1:5,000 | T2200 | Sigma-Aldrich |
| Actin | Rabbit | 1:2,000 | A2066 | Sigma-Aldrich |
| Anti-rabbit HRP | Goat | 1:50,000 | 7074 | Cell Signaling |
| DRP1 | Rabbit | 1:1,000 | 8570 | Cell Signaling |
| GBF1 | Rabbit | 1:1,000 | ab86071 | Abcam |
| Mitofusin 2 | Rabbit | 1:1,000 | 9482 | Cell Signaling |
| OPA1 | Rabbit | 1:1,000 | 80471 | Cell Signaling |
| Presenilin 1 | Rabbit | 1:1,000 | 5643 | Cell Signaling |
| Presenilin 2 | Rabbit | 1:1,000 | 9979S | Cell Signaling |
